# Cryo-EM structures reveal multiple stages of bacterial outer membrane protein folding

**DOI:** 10.1101/2021.08.24.457559

**Authors:** Matthew Thomas Doyle, John R. Jimah, Jenny E. Hinshaw, Harris D. Bernstein

**Affiliations:** Genetics and Biochemistry Branch, National Institute of Diabetes and Digestive and Kidney Diseases, National Institutes of Health, Bethesda, MD 20892 USA; Laboratory of Cell and Molecular Biology, National Institute of Diabetes and Digestive and Kidney Diseases, National Institutes of Health, Bethesda, MD 20892 USA

**Keywords:** membrane protein folding, membrane dynamics, outer membrane protein, BAM, β-barrel

## Abstract

Transmembrane β-barrel proteins are folded into the <u>o</u>uter <u>m</u>embrane (OM) of Gram-negative bacteria by the β-<u>b</u>arrel <u>a</u>ssembly <u>m</u>achine (BAM) via an unexplained process that occurs without known external energy sources. Here we used single-particle cryo-EM to visualize the folding dynamics of a model β-barrel protein (EspP) by BAM. We found that BAM binds the highly conserved “β-signal” motif of EspP to correctly orient β-strands in the OM during folding. We also found that the folding of EspP proceeds via remarkable “hybrid-barrel” intermediates in which membrane integrated β-sheets are attached to the essential BAM subunit, BamA. The structures show an unprecedented deflection of the membrane surrounding the EspP intermediates and suggest that β-sheets progressively fold towards BamA to form a β-barrel. Along with *in vivo* experiments that tracked β-barrel folding while the OM tension was modified, our results support a model in which BAM harnesses OM elasticity to accelerate β-barrel folding.

## INTRODUCTION

The insertion and folding of integral membrane proteins involves fundamentally complex processes that require management of hydrophobic and hydrophilic interfaces during intermediate steps to arrive at a topologically correct and functional structure. The biogenesis of proteins located in the outer membrane (OM) of Gram-negative bacteria and organelles of bacterial origin that span the membrane via a unique “β-barrel” structure is especially enigmatic, in part because they rapidly insert into the OM in the absence of known external energy sources (Horne et al., 2020; Tomasek and Kahne, 2021). For unknown reasons, almost all bacterial outer membrane proteins (OMPs) contain a transmembrane β-barrel. Although they can vary greatly in size (8 – 36 β-strands) and can be linked to soluble domains, transmembrane β-barrels generally conform to common architectural rules (Gruss et al., 2013; Horne et al., 2020; Lauber et al., 2018; Schulz, 2000). OMP β-barrels are tilted amphipathic anti-parallel β-sheets that are closed by tight hydrogen-bonding between the first and last β-strands (the “β-seam”) into super-stable cylinders (Horne et al., 2020; Schulz, 2000). Transmembrane β-barrels are also stabilized in the OM by parallel “girdles” of membrane-facing aromatic residues (Schulz, 2000). The majority (77%) of β-barrels also have a highly conserved C-terminal motif called the “β-signal” that contains an essential terminal phenylalanine residue of unknown function (Struyve et al., 1991; Wang et al., 2021). In bacteria, the assembly (folding and integration) of β-<u>b</u>arrels is catalyzed by a heterooligomer called the β-<u>b</u>arrel <u>a</u>ssembly <u>m</u>achine (BAM) (Heinz and Lithgow, 2014; Heinz et al., 2015; Wu et al., 2005). In *E. coli*, BAM is composed of an essential subunit (BamA), and four lipoproteins (BamBCDE) (Wu et al., 2005). BamA is conserved in all Gram-negative bacteria, and essential homologs are also found in mitochondria and chloroplasts (Heinz and Lithgow, 2014; Kozjak et al., 2003; Patel et al., 2008; Voulhoux et al., 2003). BamD is likewise highly conserved throughout bacteria but is conditionally essential (Anwari et al., 2012; Hart et al., 2020; Hart and Silhavy, 2020). BamA is itself an OMP that contains a C-terminal β-barrel domain and five soluble N-terminal <u>po</u>lypeptide <u>tr</u>ansport-<u>a</u>ssociated (POTRA) domains that bind the lipoproteins (Gu et al., 2016; Han et al., 2016; Iadanza et al., 2016).

The structural dynamics that occur as OMPs transition from an incompletely folded state to a fully folded β-barrel remain unclear. However, available evidence suggests that OMP β-signals may be recognized by BAM and that the unusual conformational malleability of BAM (particularly BamA) may facilitate the folding process (Doerner and Sousa, 2017; Doyle and Bernstein, 2019; Hagan et al., 2015; Iadanza et al., 2016; Kaur et al., 2021; Lundquist et al., 2018; Noinaj et al., 2014; Tomasek et al., 2020; White et al., 2021). Interestingly, BamA does not contain a canonical β-signal at its C-terminus, but instead has a “kinked” structure that causes its terminal residues to move dynamically and generate a unique unstable β-seam (that forms hydrogen-bonds poorly) (Lundquist et al., 2018; Noinaj et al., 2013). BamA can also adopt either inward-open or outward-open conformational states (in which the BamA β-barrel lumen is open to the inside of the cell but closed on the surface or *vice versa*) that coincide with the opening and closing of its β-seam (Gu et al., 2016). Experiments in which the BamA β-seam was tethered closed by disulfide bonds indicates that the opening and/or closing of BamA is required for efficient β-barrel folding (Gu et al., 2016; Iadanza et al., 2016; Noinaj et al., 2014). To explain the requirement for BamA β-seam opening, we recently performed an *in vivo* crosslinking study that captured a snapshot of the folding process in which the β-signal strand of an incompletely folded β-barrel was fully paired with BamA β-strand 1 (β1) via an antiparallel inter-strand interface to form a remarkable “hybrid-barrel” intermediate folding state (Doyle and Bernstein, 2019). In that study, the opposing interface between the C-terminus of BamA and the N-terminus of the β-barrel substrate was extremely dynamic, which suggests the presence of multiple transition states during the assembly process (Doyle and Bernstein, 2019). A 4 Å resolution cryo-electron microscopy (cryo-EM) structure of BAM engaged during the folding of an assembly deficient BamA deletion mutant (BamA_ΔL1_) in detergent micelles also showed BamAβ1 bound to the C-terminus of BamA_ΔL1_ to form a late-stage hybrid-barrel intermediate (Tomasek et al., 2020). Although the structure might depict a similar stage of β-barrel folding, the BAM-BamA_ΔL1_ interface is twisted and results in a “W-shaped” structure that is not fully hybridized (Tomasek et al., 2020). Due to its non-canonical final structure, it is likely that this transition state is specific to the assembly of BamA and does not occur during the folding of typical OMPs. Moreover, the BAM-BamA_ΔL1_ structure did not show how BAM recognizes the terminal phenylalanine in β-signals or reveal the dynamics of the folding process that results in the late hybrid-barrel state.

Mostly because the reconstitution of the native OM *in vitro* remains a significant technical challenge, the role of the membrane itself in OMP folding has often been neglected. Unlike other biological membranes, the bacterial OM is an asymmetric bilayer that is composed of a unique glycolipid known as lipopolysaccharide (LPS) in the outer leaflet and phospholipids in the inner leaflet (Horne et al., 2020). The concentration of OMPs within the OM is also extremely high and has been estimated to account for the majority of the OM volume (Horne et al., 2020; Jaroslawski et al., 2009). Because the interactions between densely packed β-barrels and LPS molecules results in a rigid structure in which protein diffusion is low (Rassam et al., 2015; Rojas et al., 2018; Ursell et al., 2012), the mechanism by which β-barrels are folded into the OM is even more puzzling. A recent study showed that the BAM lipoproteins can alter membrane fluidity (albeit in synthetic bilayers) and thereby potentially facilitate β-barrel integration (White et al., 2021). Intriguing molecular dynamics simulations have also raised the possibility that the unique ‘wedge-shaped” aromatic girdles of the BamA β-barrel might thin the OM to reduce the energy required for assembly (Liu and Gumbart, 2020; Noinaj et al., 2013).

Here, we examined the folding of a model *E. coli* O157:H7 OMP (EspP) that contains a stably closed β-seam, an average sized β-barrel (12 β-strands), and a canonical β-signal (Barnard et al., 2007; Franklin et al., 2018; Wang et al., 2021). By using single-particle cryo-EM to analyze an assembly-arrested form of the protein associated with BAM in native-nanodiscs that contain components directly extracted from the bacterial OM (unlike previous structural studies that analyzed BAM in detergent or nanodiscs with synthetic phospholipid bilayers), we were able to visualize multiple intermediate stages of β-barrel folding. Unlike BamA_ΔL1_, EspP forms an intermediate structure in which its conserved β-signal is fully hybridized with BamA to form a “B-shaped” hybrid-barrel. The critical phenylalanine residue in the EspP β-signal is positioned on BAM within an unusual binding pocket that interfaces with the OM to correctly orient the new OMP during folding. We also obtained direct evidence that BAM alters the structure of the OM via membrane thinning and interfacial LPS / lipid stabilization. Remarkably, in some of the intermediate hybrid-barrel structures, the OM around the folding EspP β-barrel was deflected at an angle relative to the plane of the OM around BamA. This phenomenon is unlike any known membrane-bending process (Prinz and Hinshaw, 2009). Our structural data, combined with the results of *in vivo* experiments in which β-barrel assembly was monitored during transient modulation of OM tension, led us to a completely novel model in which BAM utilizes the intrinsic structure of β-barrels and the mechanical properties of the OM itself to accelerate the final stages of OMP folding.

## RESULTS

### Structure of BAM folding a β-barrel substrate in native OM nanodiscs

To isolate an active form of BAM that is engaged in catalyzing the folding of a new β-barrel, we utilized a derivative of EspP (^MBP-76^EspP) whose assembly is arrested at a late stage while it is still bound to BAM (Doyle and Bernstein, 2019, 2021). EspP is a member of the autotransporter family of OMPs that consist of a C-terminal β-barrel and an N-terminal extracellular (“passenger”) domain that is translocated across the OM by BamA (Doyle and Bernstein, 2021; Rossiter et al., 2011). To construct ^MBP-76^EspP, we replaced most of the passenger domain with maltose binding protein (MBP), a protein that folds rapidly in the periplasm (the space between the inner membrane and OM) and, consequently, prevents translocation due to the size constraints of the channel (Doyle and Bernstein, 2019). Because translocation must be completed before BamA releases a fully folded EspP β-barrel (Ieva and Bernstein, 2009; Ieva et al., 2011), ^MBP-76^EspP remains bound to BamA in a hybrid-barrel state in which the β-signal is fully hybridized to BamAβ1 (Doyle and Bernstein, 2019). Importantly, BAM-^MBP-76^EspP co-complexes represent *bona fide* folding intermediates because β-barrel folding can be completed when the MBP containing portion of ^MBP-76^EspP is released by proteolysis (Doyle and Bernstein, 2019, 2021). To increase stability during purification, we used an *E. coli* strain transformed with plasmids expressing ^His^BamA_S425C_BCDE and ^MBP-76^EspP_S1299C_ and generated a disulfide-tether between two residues in BamAβ1 and the EspP β-signal that were previously shown to be proximal during the natural hybrid-barrel assembly step *in vivo* (Doyle and Bernstein, 2019). To more faithfully reconstitute an OM environment than previous structural studies on BAM (or other OMPs), we used a detergent-free system involving styrene–maleic acid (SMA) copolymers to directly solubilize and isolate BAM-^MBP-76^EspP co-complexes into native nanodiscs. Based on structural studies on α-helical membrane proteins, our BAM-^MBP-76^EspP OM-nanodiscs likely contain locally derived phospholipids and LPS (Lee et al., 2016; Sun et al., 2018). Purified BAM-^MBP-76^EspP OM-nanodiscs contained lipoproteins in the correct stoichiometry (Figure 1A). The BamA-^MBP-76^EspP hybrid-barrel exhibited an intrinsic feature of β-barrels when examined by SDS-PAGE in that in the absence of heat it was resistant to unfolding by SDS and migrated more rapidly (Doyle and Bernstein, 2019; Noinaj et al., 2015). Furthermore, the rapidly migrating BamA-^MBP-76^EspP hybrid-barrels also ran as diffuse bands (Figure 1A, left lane) that presumably resulted from dynamic interactions between the EspP β-barrel N-terminal strand and BamA C-terminal strands that were previously observed during folding *in vivo* (Doyle and Bernstein, 2019).

**Figure 1:**
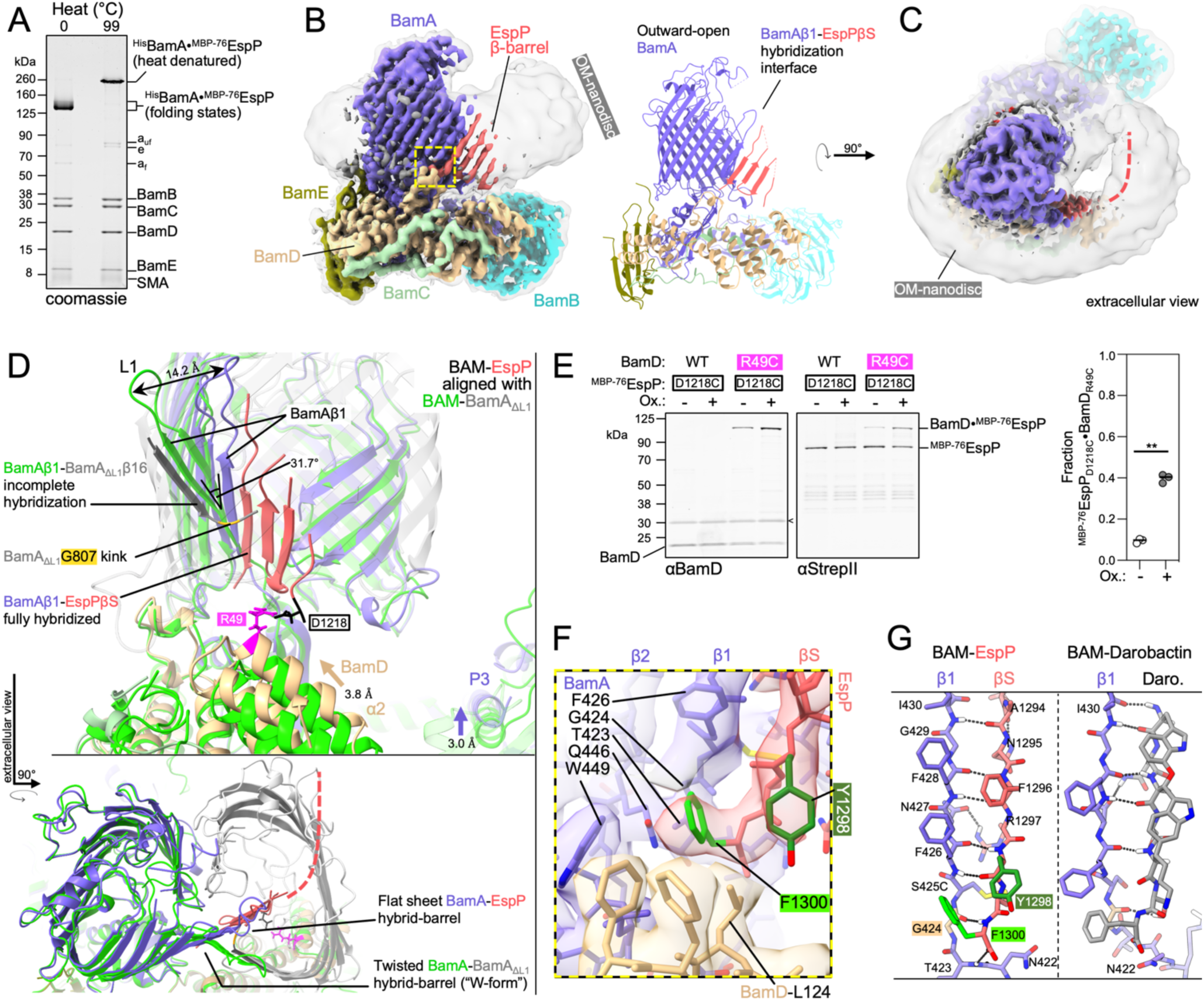
BAM binds the conserved OMP β-signal to form a flat hybrid β-sheet in the OM. **(A)** Heat denatured (99 °C) or unheated BAM-^MBP-76^EspP OM-nanodiscs were resolved by cold-SDS-PAGE. BamA and ^MBP-76^EspP are disulfide-crosslinked (•) (via S425C and S1299C, respectively). Unoxidized ^MBP-76^EspP (e), BamA folded (a_f_) and unfolded (a_uf_) species, and SMA copolymers are indicated. **(B)** High-resolution cryo-EM map (left, 3.6 Å average) and model of BAM-^MBP-76^EspP (3.6 Å average). Map colored by subunit. Local resolution filtered map at a lower threshold level (left, transparent grey) shows OM disc boundary. BamA is in an outward-open conformation and hybridized to the EspP β-signal (βS) strand via BamA β-strand 1 (β1). Yellow box shown in **F**. **(C)** Extracellular top view of map as in **B**. Dashed line indicates likely location of remainder of EspP β-barrel. **(D)** Substrate-specific intermediate states and BAM conformations during the assembly of EspP β-barrel and BamA_ΔL1._ Structure of BAM-BamA_ΔL1_ hybrid-barrel intermediate state in detergent (PBD 6V05) (BAM, green; BamA_ΔL1_, transparent grey) is aligned with the BAM-^MBP-76^EspP structure (subunit colors as in **C**). The noncanonical final strand of BamA_ΔL1_ is not fully hybridized with BamAβ1 (kink at BamA_G807_, yellow) whereas the conserved EspP β-signal strand is fully hybridized with BamAβ1. A 31.7° difference in BamAβ1 tilt angle [axis residues 427 – 434 alpha carbons (αC)] coincides with either “flat-sheet” (BamA-EspP) or twisted “W-shaped” (BamA-BamA_ΔL1_) hybrid-barrel assembly intermediates. BamA POTRA3 (“P3”) and BamD α-helix 2 (“α2”) are denoted. **(E)** *E. coli* BL21(DE3) expressing ^His^BamABCDE (or ^His^BamABCD_R49C_E) and ^MBP-76^EspP_D1218C_ (residues mutated to cysteine indicated in **D**) were mock treated (Ox. -) or treated with 4-DPS (Ox. +) and BamD•^MBP-76^EspP crosslinks were identified by double-immunoblotting with αBamD and αStrepII (StrepII-tag at ^MBP-76^EspP N-terminus) antibodies (n = 3). Non-specific band is denoted (<). The graph (right) shows the fraction of crosslinked BamD•^MBP-76^EspP [line at median, two-tailed paired t-test: *P* = 0.0019 (**)]. **(F)** β-signal terminal residue binding pocket (magnified yellow box in **B**). Highly conserved Y(−3, dark green) and F(−1, light green) β-signal residues are indicated. **(G)** Comparison of BamAβ1-EspP β-signal strand and BamAβ1-darobactin (PDB 7NRI) interactions. In both cases F(−1) is positioned over the space created by BamA_G424_ (tan). Dotted lines denote H-bonds.

A high-resolution structure of the purified BAM-^MBP-76^EspP OM-nanodiscs was solved to a global resolution of 3.6 Å by single particle cryo-EM (Figure 1B & Figure S1). The structure revealed BamA in an overall outward-open conformation with BamAβ1 associated with the EspP β-signal to form a hybrid-barrel intermediate folding state (Figure 1B). In this map only four C-terminal β-strands of the actively folding EspP β-barrel were clearly resolved. These β-strands extended into a low-resolution region within the OM-nanodisc that likely represents the remainder of the amphipathic EspP β-barrel creating a border between the membrane and an internal hydrophilic cavity (Figure 1C). The low resolution of the EspP β-barrel N-terminus suggests that this portion of the protein transitions between multiple highly dynamic folding substates, a notion consistent with the previously observed dynamic interface between the EspP β-barrel N-terminus and BamAβ15/16 mentioned above (Doyle and Bernstein, 2019).

Comparison of our structure to the BAM-BamA_ΔL1_ structure (Tomasek and Kahne, 2021) showed striking differences. While the hybridization interface between BAM and the BamA_ΔL1_ mutant is twisted to form a W-shaped hybrid-barrel, the BamA-EspP hybridization interface instead forms a continuous flat β-sheet (Figure 1D). This difference stems from the ability of BamAβ1/2 to flex and tilt in the membrane and suggests a mechanism by which BamA can accommodate the folding and integration of different β-barrel substrates. In the BAM-^MBP-76^EspP structure, BamA POTRA3 and BamB (through its association with POTRA3) are also positioned closer to the membrane (Figure 1D “P3” & Figure S1). Conformational changes localized near the N-terminal α-helices of BamD likewise result in a shift towards the membrane with α-helix 2 interfacing with the outer side of the periplasmic turns of the folding EspP β-barrel (Figure 1D & Figure S1). This overall conformation contrasts with the BAM-BamA_ΔL1_ structure in which BamD is positioned beneath the lumen of BamA_ΔL1_. To test whether BamD can interact with the periplasmic turns of EspP during a hybrid-barrel stage of assembly *in vivo*, BAM containing a BamD_R49C_ subunit was co-expressed in *E. coli* with ^MBP-76^EspP_D1218C_ (cysteine substitution sites are indicated in Figure 1D) and cells were treated with a thiol-specific disulfide-oxidation catalyst. Consistent with our structure, strong disulfide-crosslinking between ^MBP-76^EspP_D1218C_ and BamD_R49C_ was observed after chemical oxidation but not in the control strain expressing a wild type (WT) BamD allele (Figure 1E). The observation that crosslinking plateaued at ∼40% suggests the presence of additional substates with alternative conformations of EspP relative to BamD (Figure 1E plot & Figure S1). A higher than expected level of spontaneous crosslinking (∼10%, Figure 1E, Ox-) also indicated the existence of a stage in which EspP interacts with BamD very stably.

Strikingly, in the BAM-^MBP-76^EspP structure we were able to clearly resolve the conserved residues of the canonical β-signal of EspP (Figure 1B & 1F). The terminal EspP β-signal residue (F1300) is oriented over BamAβ1 in a space created by BamA_G424_ that forms a novel structural arrangement reminiscent of stabilizing intra-barrel “mortise-tenon joints” (Figure 1F &1G) (Leyton et al., 2014). BAM interacts with F1300 via BamA T423, G424, F426, and Q446 within an unusual membrane facing hydrophobic pocket (Figure 1F). Nevertheless, the β-signal binding pocket is not totally filled. Presumably this property enables BAM to accommodate the less common subset of OMPs that have β-signals terminated by tryptophan or tyrosine instead of phenylalanine (Struyve et al., 1991; Wang et al., 2021). The conserved β-signal residue at the -3 position of EspP (Y1298) interacts with BamD L124 and is oriented into the membrane plane at a depth corresponding to the aromatic girdles of fully folded canonical β-barrels (Figure 1F). Overall, the structure suggests that BAM binds to β-signals to correctly orient the C-terminal strands of new OMPs into the OM during the folding process. Interestingly, darobactin (a recently discovered BAM inhibitor) (Imai et al., 2019) and the β-signal interact with BamAβ1 in a highly similar way; like EspP F1300, the terminal phenylalanine of the darobactin peptide is positioned over BamA_G424_ (Kaur et al., 2021) (Figure 1G). Therefore, our structure not only provides the structural basis for native OMP β-signal binding by BAM during assembly, but also definitively shows that darobactin is a competitive inhibitor of OMP substrate recognition and thereby helps to explain its bactericidal potency.

An important aspect of our study is that by solving the structure of BAM-^MBP-76^EspP within native nanodiscs that harbor local OM lipids captured during solubilization, we can consider the role of the of the OM in OMP assembly. In our BAM-^MBP-76^EspP map, we observed a repetitive pattern of stabilized density circling the BamA β-barrel at the expected location and size of outer leaflet LPS lipid A head groups and clear boundaries that likely represent density for inner leaflet phospholipid headgroups (Figure 2A left & middle). It has been postulated that BamA locally thins the OM to decrease the energetic penalty of OMP integration (Liu and Gumbart, 2020; Noinaj et al., 2013). To test this hypothesis, we measured the membrane thickness at positions that had clear and repeatable density proximal to BamA (Figure 2A right & Figure S2). The membrane near the N-terminal half of the BamA β-barrel (∼β1 – β7) is ∼25 – 26 Å in width. This value corresponds closely to the estimated average thickness of the OM (∼25 Å) (Wu et al., 2014) and is slightly thicker than the average hydrophobic region of OMPs (∼24 Å) (Lomize et al., 2011). Interestingly, the side of the OM-nanodiscs near strands β8 – β13 exhibited prominent local thinning to a “pinch-point” (∼20 Å) that thickens again near the C-terminal BamA curl. These observed membrane depth patterns perfectly match recent molecular dynamics simulations that predict thickening/thinning patterns around BamA (Liu and Gumbart, 2020). The map also suggests even more extreme membrane thinning across strands β14 – 16, but the density does not have clear boundaries for measurement. At the location of the pinch-point we also observed striking density that likely corresponds to a lipid A moiety with a single stabilized acyl chain (the other lipid A acyl chains are presumably dynamic) (Figure 2B). The stabilized acyl chain lies within a groove alongside BamA strands β 11/12 that is created by the lipid-facing residues G631 and A714. Because the membrane thickness and stabilization patterns observed in the high-resolution BAM-^MBP-76^EspP structure were likewise observed in our subsequent independent BAM-^MBP-76^EspP reconstructions (see below and Figure S2), they are likely valid structural features. We speculate that the stabilization of lipid acyl chains on the C-terminal side of BamA is helpful for the process of membrane thinning.

**Figure 2:**
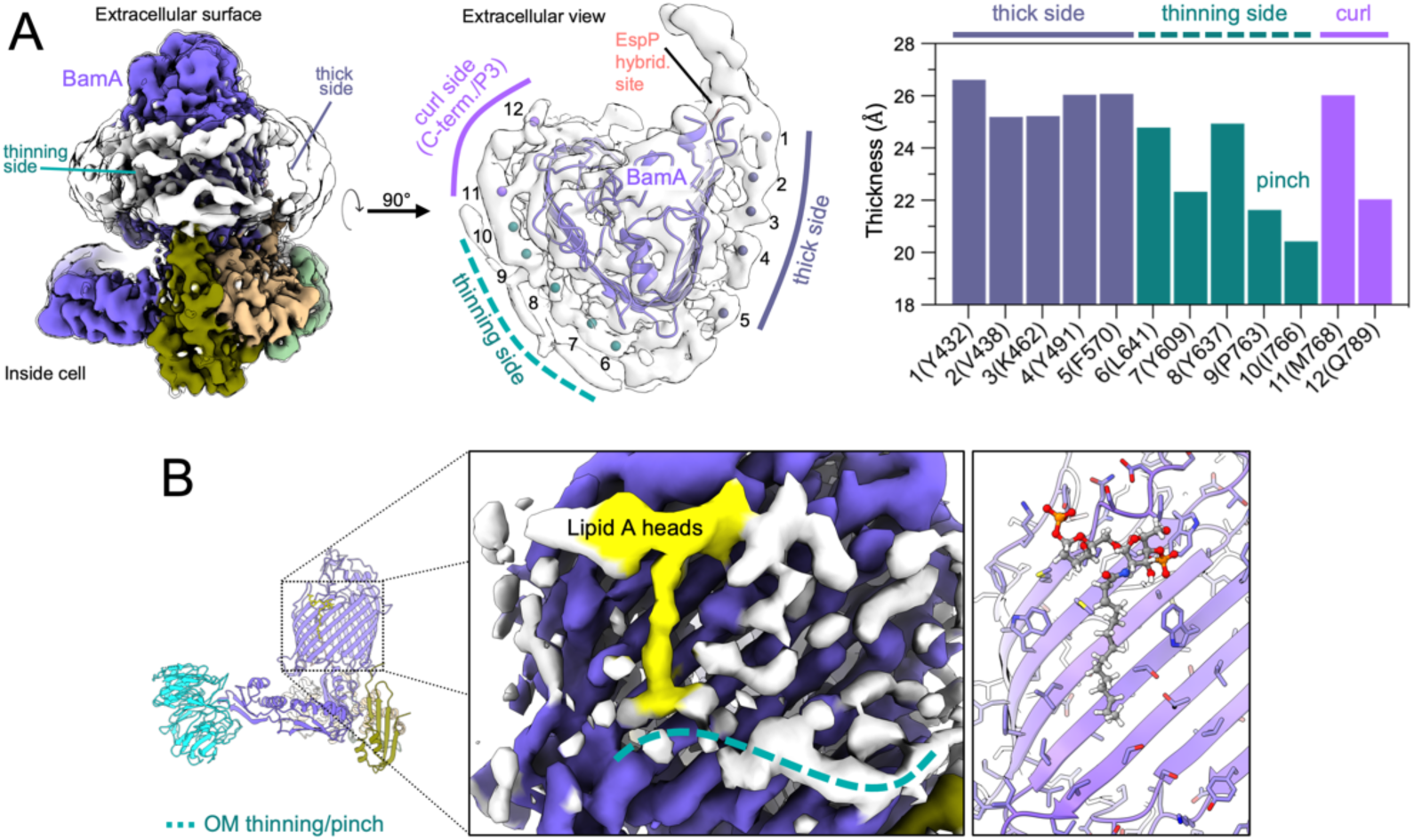
OM thinning and lipid/LPS ordering during OMP folding. **(A)** Left, side view of BAM-^MBP-76^EspP showing both thin and thick membrane sides around BamA [local resolution filtered map to showing protein (colored) and membrane (white) features and at a lower threshold (clear) to show OM-disc boundary]. Middle, extracellular view of map with measurement marker positions (circles) indicated. Repeated pattern in the OM-disc indicative of outer leaflet interfacial lipid A head groups. Right, thickness measurements of membrane density surrounding the BamA β-barrel at indicated positions (residues close to outer leaflet markers in brackets). **(B)** BAM-^MBP-76^EspP map shows density consistent with a lipid A head groups and a stabilized acyl chain (yellow) (modeled on right) on the thinning side of the BamA β-barrel (teal dashed line).

### The BAM, the OM, and the incoming OMP each undergo major structural transitions during β-barrel folding

To generate the BAM-^MBP-76^EspP structure described in the preceding section, we started from a pool of ∼1.2M high quality particles generated in RELION and obtained the high-resolution map after rounds of heterogenous refinement in cryoSPARC. Although our map had a higher global resolution than the previously solved BAM-BamA_ΔL1_ structure, the local resolution was poor in the area corresponding to the N-terminal portion of the EspP β-barrel and low for BamA P3 / BamB and the N-terminus of BamD (Figure 3A) presumably due to significant dynamicity in these regions. During processing we noticed specific low-resolution classes that appeared to have alternate conformations in these areas and wondered whether a more conservative processing strategy could improve the local resolution, albeit at the expense of global resolution. To that end, we reprocessed the entire ∼1.2M particle pool in RELION into 6 classes and then separately processed each class in cryoSPARC yielding reconstructions with global resolutions between 4.2 – 4.5 Å (Figure 3B & Figure S3). The conformation of the BamA β-barrel is essentially identical in all of the structures (Figure 3C). Classes 3, 4 and 6 are similar to the original high-resolution structure but contained slight changes in the position of BamB and the N-terminus of BamD. The density of BamB is poor in class 2 (although it is visible at lower thresholds) presumably because it is highly dynamic (Figure 3B). Indeed, between all the classes the largest overall BAM conformational changes were in the positioning of BamB (Figure 3C & Video S1). Classes 1 and 5 represent conformational extremes in which the BamA POTRA domains and BamD move closer or farther away from each other and, concomitantly, BamA P3 moves towards or away from the membrane (Figure 3D). Consequently, this results in very significant changes in BamB positioning (Figure 3D). Together, these reconstructions show that BAM periplasmic components undergo large conformational changes during the late stages of OMP folding that mimic the structural heterogeneity observed in the periplasmic subunits even in apo-BAM (Iadanza et al., 2020).

**Figure 3:**
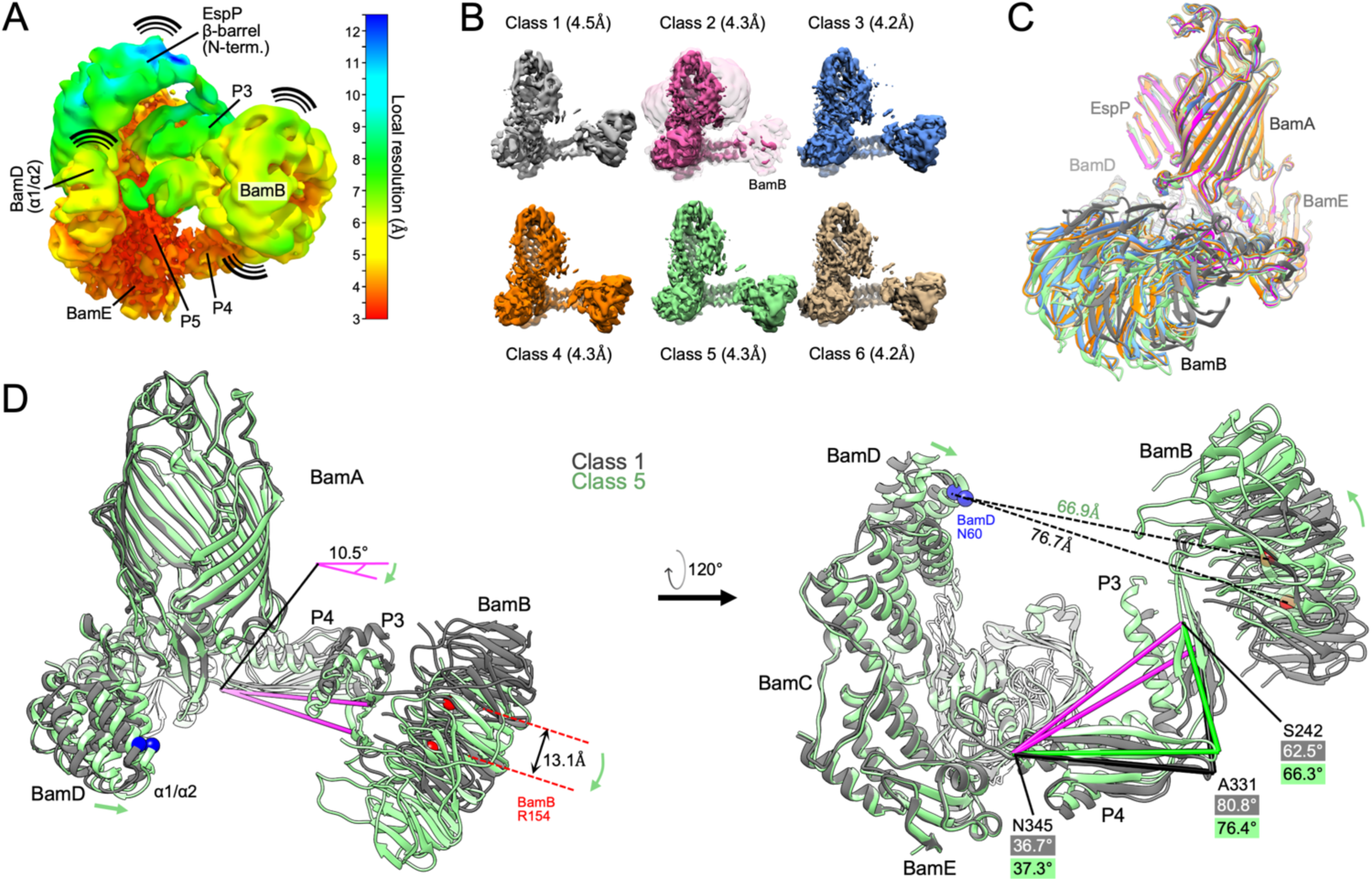
Conformational changes of BAM periplasmic components during OMP folding. **(A)** The BAM-^MBP-76^EspP high-resolution map (shown in Figure 1B) filtered and colored by local resolution. Low resolution components [e.g. EspP β-barrel N-terminus, BamB, and POTRA3 (P3)] are conformationally dynamic. High-resolution components (e.g. BamE and P5 are conformationally stable. **(B)** Conformationally diverse cryo-EM maps of BAM-^MBP-76^EspP. Maps filtered by local resolution. Class 2 is shown at a lower threshold level (clear pink) to show BamB **(C)** Models of BAM-^MBP-76^EspP Classes 1 – 6 were aligned (based on BamA P5 residues Y348 – R421). View shows large conformational variability in BamB positioning. Colors as in **B**. **(D)** Classes 1 and 5 aligned as in **C**. Conformational changes are depicted by green arrows. Axis (pink) from BamA S242 (P3) to N345 (hinge region between P4 & P5) α-carbons flexes by 10.5° between classes. Changes in the angles between S242, A331, and N345 also shows flexing between P3 & P4 between classes.

The simple classification approach did not improve the maps in the region corresponding to the folding EspP β-barrel. To better resolve the intermediate folding states of EspP, we subtracted the signals for BamB, BamA P3, and the BamD N-terminus from the original ∼1.2M particle pool and conducted focused 3D classification and refinement on the remaining complex. We reasoned that removing these dominant sources of structural heterogeneity would allow better alignment of the conformational substates of the EspP β-barrel N-terminus during the hybrid-barrel stage that were predicted from our earlier experiments *in vivo* (Doyle and Bernstein, 2019). Consistent with our hypothesis, we were able to generate multiple reconstructions of novel hybrid-barrel substates using this processing strategy (Figure 4 and S3). In one structure that we designate the “<u>o</u>pen-<u>s</u>heet” (OS)-state (Figure 4A), the EspP β-barrel is observed as a remarkable membrane-integrated open β-sheet with its C-terminus hybridized to BamA. In this state the OM-nanodisc is deflected around the EspP transmembrane β-sheet at an angle that results in a mismatch of the membrane plane around BamA (Figure 4A). In the “intermediate-<u>o</u>pen” (IO)-state (Figure 4B), the reconstructed BAM components are essentially identical to the OS-state with both structures showing BamA in an outward-open conformation. However, compared to the OS-state, in the IO-state the EspP transmembrane β-sheet is folded closer to BamA and the observed membrane deflection is less extreme. In a third structure that we call the “<u>b</u>arrelized/<u>c</u>ontinuous-<u>o</u>pen” (B/CO)-state, we observed a “B-shaped” BamA-EspP hybrid-barrel but, unlike the other states, no obvious membrane deflection (Figure 4C). This state presumably represents a very late stage of EspP assembly in which the β-barrel structure is nearly complete. In the B/CO-state, BAM is observed in a totally novel conformation in which the C-terminal half of the BamA β-barrel is expanded away from the N-terminus and repositioned higher in the membrane plane so that its surface loops (including L4, 6, and 7) are shifted away from the EspP β-barrel (Figure 4D). The result is a BamA structure reminiscent of outward-open states but with an opening that creates a continuous channel through the OM-nanodisc (Figure 4D). This structure may represent a stage prior to the release of the EspP β-barrel that we have recently observed *in vivo* in which the BamA β-barrel facilitates secretion of the EspP passenger domain (Doyle and Bernstein, 2021).

**Figure 4:**
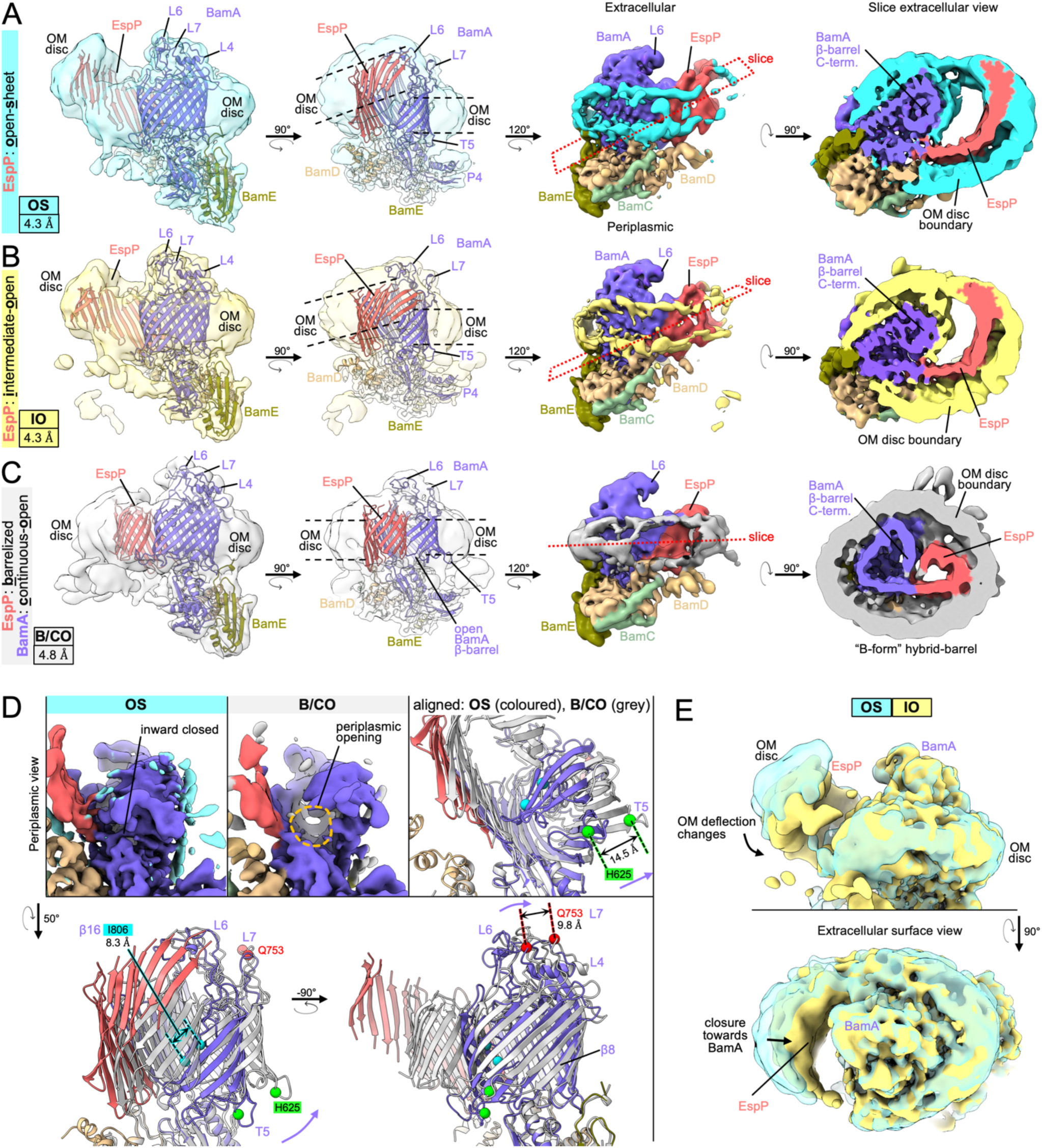
Focused classification reveals a continuous-open BamA β-barrel and multiple hybrid-barrel substates during EspP folding. **(A-C)** Focused classification and refinements on particles with subtracted signals of dynamic periplasmic components (BamD N-term., P3, and BamB) identified three distinct BamA-EspP hybrid-barrel conformations: EspP “open-sheet” (OS, panel **A**, OM-disc in cyan), EspP “intermediate-open” (IO, panel **B**, OM-disc in yellow), and EspP “barrelized” / BamA “continuous-open” (B/CO, panel **C**, OM-disc in light gray). Map resolutions are quoted at bottom left. Dashed lines in mid-left views indicate in-plane or deflected bilayer angles. Right views, maps are colored by subunit. Far-right shows slices across the BamA-EspP hybrid-barrels (slice planes depicted mid-right) with expanded hybrid-barrels for OS and IO, and a “B-shaped” hybrid-barrel for the B/CO state. Maps are filtered by local resolution. **(D)** Identification of a novel BamA conformation during OMP assembly. Compared to the OS class (and all other reconstructions in this work) the B/CO class exhibits the BamA β-barrel in an expanded conformation similar to outward-open conformations but with a periplasmic opening (compare maps top left and middle, periplasmic view). In top-right and bottom panels, models of OS and B/CO states are aligned (on P5 residues Y348 – R421) showing that the C-terminal half of the BamA β-barrel both expands and shifts towards the cell surface through changes beginning at strand β8. The expansion also coincides with extracellular loops (L4, 6, and 7) moving away from the hybridization interface. Differences in selected residue αC positions in turn 5 (T5, H625), L7 (Q753), and β16 (I806) between OS and B/CO states are denoted. **(E)** Overlay of OS and IO maps (filtered by local resolution) shows changes in the degree of OM-bilayer deflection and degree of closure in the EspP β-sheet region.

### Antagonism between the intrinsic structure of OMPs and OM tension drives late folding

Although we cannot definitively order the three hybrid-barrel substates in a temporal sequence because they are derived from the same sample, a simple interpretation of the data is that the OS-state represents an early stage of folding following the membrane integration of the EspP β-sheet, the IO-state represents a slightly later stage in which the EspP β-sheet folds towards BamA (Figure 4E), and the B/CO-state represents a relatively late stage in which the β-sheet folds into a barrel-like structure (see Video S2). It is notable that the changes in EspP folding between OS and IO states were not associated with major structural changes in BamA. Furthermore, the extreme nature of the expanded EspP β-sheet and membrane deflections in the OS and IO-states were very surprising and warranted an explanation that places these states in the context of folding within the native OM.

To rationalize our findings, we conceptualized a new model of the late stages of OMP assembly by considering the intrinsic structure of transmembrane β-barrels and the rigidity of the OM. OMP β-strands are tilted by ∼45° from the barrel-axis (Schulz, 2000) and, due to the presence of aromatic girdles and other membrane-facing hydrophobic residues in β-barrel transmembrane β-sheets, fully folded OMPs are often slightly tilted in the OM (Lomize et al., 2012). We calculated the membrane orientations of solved *E. coli* β-barrels and then illustrated them as open β-sheets while maintaining their β-signals in their calculated positions relative to the OM plane (Figure 5A right, Figure S4A). The result is a mismatch in which the N-terminal transmembrane β-strands would not reside in the normal OM plane but would instead deflect the membrane. Consistent with this hypothetical scenario, the OS and IO-state structures capture EspP as an incompletely-folded open β-sheet at an angle relative to the normal OM plane (Figures 4A, 4B, 5B and S4B). Indeed, in all our structures we observe BamA bound to the EspP β-signal at an even higher angle than the β-signal is situated in after EspP is completely folded (∼64.6° vs. 58°, Figure 5A, 5B, & Figure S4B). Furthermore, the difference between the angle of the β-signal of some OMPs when bound to BamA as an open β-sheet versus the fully folded form may be even greater (Figure S4A, see PgaA). The outward-open BamA conformation might therefore create a highly antagonistic scenario between the normal OM plane and the intrinsic structure of incompletely folded β-barrels that deflects the membrane (Figure 4A, 4B, 5B). This scenario, however, should be considered in light of the finding that unlike other biological membranes, the OM forms a very rigid structure (stiffer than the peptidoglycan cell wall) due to interactions between LPS molecules, the high OMP density, and the stiffness of OMPs themselves (Horne et al., 2020; Jaroslawski et al., 2009; Lessen et al., 2018; Rojas et al., 2018). Indeed, due to its rigidity, the OM has been observed to function as a ‘spring’ that undergoes compressive changes during osmotic shock (Rojas et al., 2018). Although stretching and compressive forces appear to be globally equalized across the *E. coli* OM during steady-state growth (Rojas et al., 2018), our model predicts that the intrinsic structure of hybrid-barrel deflected β-sheets (Figure 4A, 4B) would be countered by the intrinsic local tensile forces of the OM to help drive the closure of β-barrels (Figure 5B).

**Figure 5:**
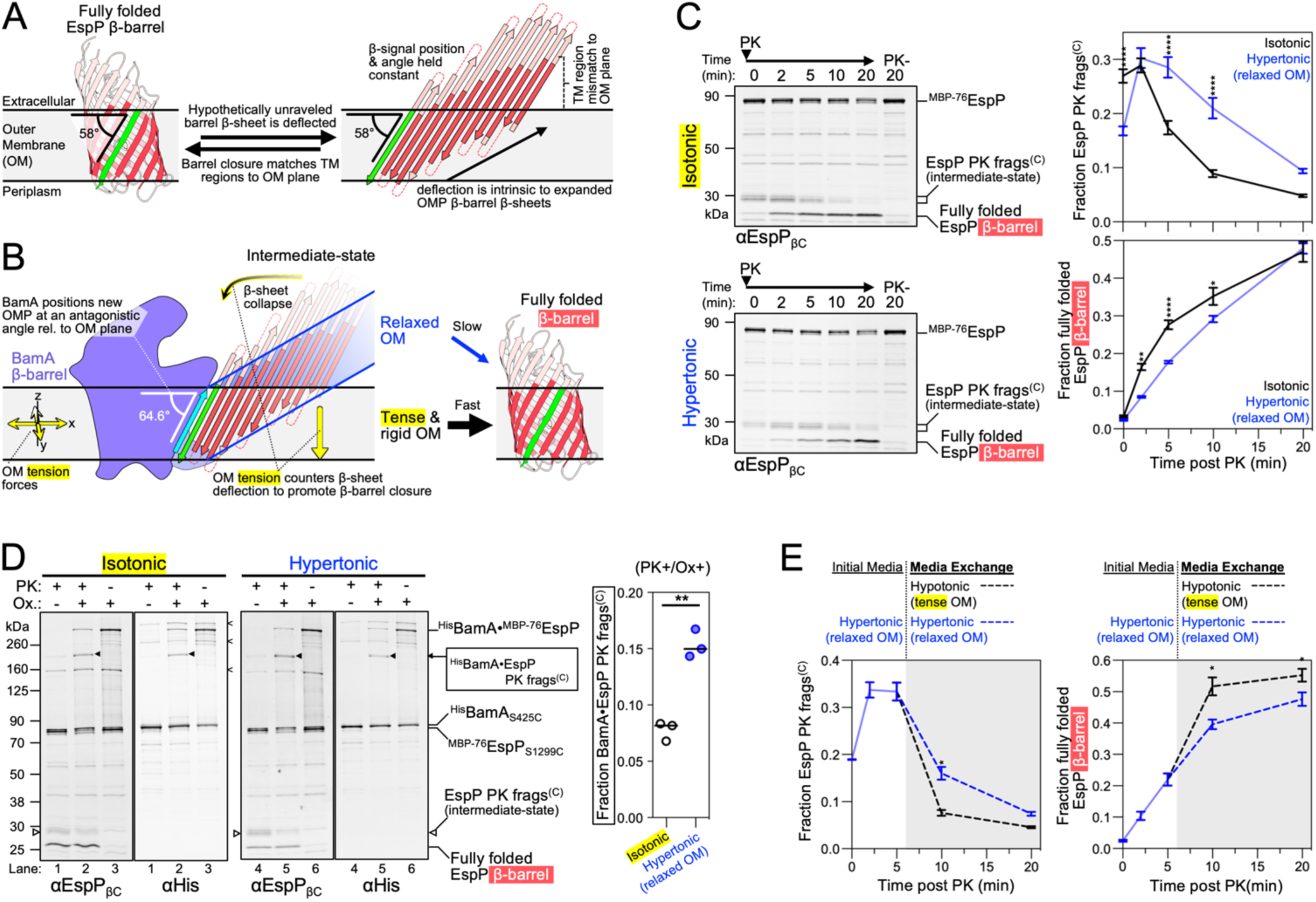
OM tension accelerates transmembrane β-barrel folding. **(A)** Hypothetically unraveled EspP β-barrel with the position of the conserved β-signal strand (green) held constant to its angular membrane orientation when fully folded (58° for EspP, membrane plane calculated using OPM server) (Lomize et al., 2012) showing intrinsic mismatch between the membrane plane and the transmembrane portion of β-strands (red). Depicted loops and turns are not drawn to-scale. **(B)** EspP β-sheet as in **A** except model depicted with experimentally determined angle (using the BAM-^MBP-76^EspP high resolution structure) of EspP β-signal relative to the membrane when hybridized to BamAβ1 (cyan). The model predicts that the intrinsic tension forces of the rigid OM of Gram-negative bacteria aids β-barrel folding by countering the intrinsic deflection of an incompletely folded expanded β-sheet. **(C)** *E. coli* BL21(DE3) expressing ^His^BamABCDE and ^MBP-76^EspP were suspended in LB (isotonic control) or LB-sorbitol (hypertonic) to generate cells with a relaxed OM. β-barrel folding was restarted from the hybrid-barrel stage by adding PK to release a C-terminal EspP fragment (frags^(C)^). Fragment conversion to a completely folded β-barrel after PK addition was monitored by immunoblotting using an antiserum generated against the C-terminus of EspP (αEspP_βC_). Left, representative blots. Right, mean fraction of the EspP PK fragment (top) or converted into a folded β-barrel (bottom) (±SEM, n = 4). **(D)** *E. coli* BL21(DE3) expressing ^His^BamA_S425C_BCDE and ^MBP-76^EspP_S1299C_ were grown and treated as in **C** except that samples at 5 min post PK were mock treated (Ox. -) or treated with 4-DPS (Ox. +) and ^His^BamA•^MBP-76^EspP crosslinks were identified by double-immunoblotting with αEspP_βC_ and αHis antibodies (n = 3). Non-specific crosslinks are denoted (<). Right, fraction of crosslinked BamA•^MBP-76^EspP C-terminal PK fragment out of all bands detected with αEspP_βC_ [line at median, two-tailed paired t-test: *P* = 0.0038 (**)]. **(E)** Bacteria were grown, suspended in LB-sorbitol, and treated with PK as in **C** except that after 5 min the media was exchanged for either LB-sorbitol (hypertonic) or LB (hypotonic) (gray plot area). Plots are mean fraction (±SEM, n = 3). For **C** and **E**, 2-way repeated measures ANOVA (Šídák’s tests) were performed *p < 0.05, **p < 0.01, and ****p < 0.0001.

To test our model *in vivo*, we expressed ^MBP-76^EspP alongside BAM in *E. coli* to create a pool of BamA-^MBP-76^EspP hybrid-barrels in the OM and then subsequently monitored late-stage β-barrel folding kinetics during modification of OM tension via osmotic shock. As mentioned earlier, the assembly of ^MBP-76^EspP can be restarted from the hybrid-barrel stage by adding a protease that removes the MBP portion responsible for arresting assembly. Completion of folding can then be assessed by monitoring the auto-catalytic cleavage of the passenger domain that occurs after it is fully secreted and the β-barrel reaches its native conformation (Dautin et al., 2007; Ieva and Bernstein, 2009). Consistent with previous studies (Doyle and Bernstein, 2019, 2021), adding proteinase K (PK) to bacteria suspended in isotonic LB medium released ∼30 kDa C-terminal EspP fragments from the fusion protein that were rapidly converted into folded ∼27 kDa β-barrels and a peptide derived from the passenger domain that was not detected (Figure 5C top-left and black curve). However, when bacteria were exchanged into a hypertonic LB medium to relax the OM (Rojas et al., 2018) prior to the addition of PK, the incompletely folded C-terminal EspP fragments accumulated and the rate of their conversion to fully folded β-barrels was significantly reduced (Figure 5C bottom-left and blue curve). Although the β-barrel assembly delay under hypertonic conditions was most notable at 5 min after PK addition, by 20 min there was no difference in the level of folded β-barrel between the two conditions (Figure 5C). This observation strongly suggests that the delay was due to an energetic effect and that the hypertonic conditions did not simply block completion of β-barrel folding. To directly pinpoint the delay to the period that follows the formation of the hybrid-barrel but that precedes the completion of EspP β-barrel folding, the experiment was repeated using the strain expressing ^His^BamA_S425C_BCDE and ^MBP-76^EspP_S1299C_. Samples were treated with PK for 5 min (or mock-treated) and oxidized to promote disulfide-crosslinking as described above. Consistent with previous results (Doyle and Bernstein, 2019), strong crosslinks between BamAβ1 and the ^MBP-76^EspP β-signal were detected in oxidized samples without PK treatment (Figure 5D, lanes 3 and 6). In samples that were both oxidized and PK treated, the incompletely folded EspP C-terminal fragments were likewise crosslinked to BamA (Figure 5D, lanes 2 and 5, black arrows). These results confirm that hypertonic conditions do not interfere with the stability of the BamA-^MBP-76^EspP assembly intermediate and pinpoint the delay to the period following the formation of the hybrid-barrel. Interestingly, EspP C-terminal fragments crosslinked to BamA at a statistically higher level under hypertonic conditions at the 5 min time-point (Figure 5D, graph). This finding suggests that the association of incompletely folded EspP with BamA at the hybrid-barrel stage is prolonged when the OM is relaxed.

Finally, our model not only predicts that folding can be slowed by relaxing the OM, but that folding can be accelerated by increasing the OM tension. To test this idea, we repeated the assembly restart experiment again under hypertonic conditions, but 5 min after the addition of PK we exchanged the bacteria into an equivalent solution (control) or a hypotonic solution (to increase OM tension) and continued to monitor EspP β-barrel folding. Consistent with our hypothesis, the EspP C-terminal fragments were converted more rapidly into folded β-barrels when they were exchanged into a hypotonic medium (Figure 5E). The difference constitutes a substantial effect given that only a small fraction of incompletely folded EspP molecules remained to be tracked after the 5 min time-point. Together, these results support a model in which BamA orients OMP substrates at an antagonistic angle to the OM to exploit the intrinsic tension of the OM as a useful driving force to accelerate β-barrel folding (Figure 5B).

## DISCUSSION

Although the first transmembrane β-barrel structure was solved in 1990 (Weiss et al., 1990), how they are recognized, folded, and released into the bacterial OM in the absence of any known external energy sources remains poorly understood. In this work we provide structural and biochemical evidence that helps to explain all of these critical stages of OMP assembly. We solved ten distinct cryo-EM structures of BAM bound to EspP, a model OMP that contains conserved features found in most transmembrane β-barrels, that likely illustrate distinct intermediate stages in the folding process. Because the co-complex was purified in native nanodiscs instead of detergent or nanodiscs containing synthetic phospholipid bilayers, we were able to obtain insight into how OM lipids contribute to the β-barrel folding process. With respect to substrate recognition, our structural data reveal an unusual pocket in BAM that binds to the highly conserved C-terminal OMP β-signal motif. During the hybrid-barrel intermediate stage, the terminal aromatic residue interacts with several adjacent BAM residues and is positioned over the space created by BamA_G424_. The interaction resembles G-(F/Y/W)-based inter-strand mortise-tenon joints that are found within most OMPs and that provide structural stability (Leyton et al., 2014). However, the BamA-β-signal interaction constitutes the first observation of an inter-barrel mortise-tenon-like joint. The binding of BAM to the β-signal provides an explanation for the finding that mutations of the terminal aromatic residue cause severe assembly defects and lead to OMP degradation *in vivo* (Gessmann et al., 2014; Lee et al., 2018; Wang et al., 2021). Presumably the mutations reduce the binding affinity of incoming OMPs to BAM, prevent their progression to the hybrid-barrel stage, and result in exposure to periplasmic proteases. Our discovery of this binding site may also enable the design of novel competitive inhibitors of β-signal binding and the further development of lead compounds such as darobactin (Imai et al., 2019; Kaur et al., 2021) that act as potent antibiotics against multidrug-resistant Gram-negative pathogens.

Our results provide evidence that the EspP C-terminal domain inserts into the OM as a β-sheet and then folds into a β-barrel in multiple steps. The structural data suggest that BAM binds to the β-signal strand of EspP to form a flat hybridization interface and that this hybrid-barrel intermediate passes through several substantially different stages of folding (e.g., OS, IO, and B/CO substates) resulting in a “B-shaped” hybrid-barrel. Ultimately, because the BamAβ1-EspP(β-signal) backbone hydrogen-bond network is weaker than that of the β-seam of fully folded EspP (Figure S4C), this configuration provides an energetically favorable mechanism for the release of the substrate into the lipid bilayer. Furthermore, we identified a substate in which BamA exists in a novel continuous-open conformation coinciding with a more barrel-like EspP structure (the B/CO-state). We recently showed that the unfolded passenger domain of ^MBP-76^EspP is secreted through the BamA β-barrel lumen during a hybrid-barrel assembly stage *in vivo* (Doyle and Bernstein, 2021). We therefore speculate that the continuous opening in BamA observed in the B/CO-state may constitute a channel for the secretion of autotransporter passenger domains and extracellular segments of other OMPs. Despite the stability of ^MBP-76^EspP, we found that the passenger domain is extremely dynamic within the channel *in vivo* (Doyle and Bernstein, 2021), and this dynamicity may explain the lack of passenger domain density within the BamA opening in our reconstructions. Alternatively, the conformational changes observed in the surface loops of BamA in the B/CO-state relative to our other structures (e.g., L4, 6, and 7) may be required for the folding of β-barrels more generally. Indeed, this finding may explain why the function of BAM is strongly inhibited when BamA L4 is bound by the bactericidal antibody fragment Fab1 which presumably prevents this conformational cycling (White et al., 2021). It is important to note that none of our reconstructions exhibited the twisted interface that results in a hybrid-barrel with a “W-shaped” cross-section observed in the BAM-BamA_ΔL1_ structure (Tomasek et al., 2020). Besides lacking a canonical C-terminal β-signal, the BamA_ΔL1_ β-barrel substrate has unique features such as an unstable β-seam, extreme structural dynamism, and a kinked C-terminus that likely causes the twisted hybridization interface observed during its assembly (Doerner and Sousa, 2017; Gu et al., 2016; Iadanza et al., 2016; Lundquist et al., 2018; Noinaj et al., 2014; Noinaj et al., 2013; Tomasek et al., 2020). Therefore, the W-form hybrid-barrels probably represent a late intermediate stage that is unique to the assembly of BamA. Because EspP follows the common architectural rules of most β-barrels, we speculate that the majority of OMPs are folded through a late B-form hybrid-barrel stage before β-signal exchange and β-seam closure causes the release of the fully folded β-barrel.

Our structural data also enable us to discriminate among a variety of models for BAM function that have been previously proposed. We found that during its assembly by BAM, EspP can associate with BamA to form structurally diverse hybrid-barrels and that in two reconstructions EspP was observed in remarkable integrated open β-sheet conformations. Based on these findings we propose a β-barrel folding model in which the open β-sheets close towards BamA and then curl inwards to form a barrel-like structure at a late stage (Video S2). The interface between BamAβ1 and the EspP β-signal does not change significantly between our ten structures, yet the N-terminus of the EspP β-barrel undergoes enormous conformational changes. These observations are fundamentally inconsistent with “threading” models which propose that unfolded OMPs enter the BamA β-barrel lumen and form β-hairpins that are sequentially integrated into the lipid bilayer through a “lateral gate” between BamA β1 and β16 (Horne et al., 2020; Tomasek and Kahne, 2021). In contrast, our structures are consistent with our previous study in which we showed that the interface between the BamA C-terminus and the EspP β-barrel N-terminus is extremely dynamic but that the BamAβ1-β-signal interface is remarkably stable during the hybrid-barrel stage *in vivo* (Doyle and Bernstein, 2019). Based on the results, we proposed that the N-terminus of OMP β-barrels undergo a swinging action in the membrane during their assembly. In light of our structural data, we speculate that 1) at early stages of folding OMP β-signals are bound by BamAβ1, 2) this interaction templates the folding of the adjacent OMP β-strands via β-augmentation (Remaut and Waksman, 2006) until an elongated β-sheet is formed and 3) during the folding process the BamA β-barrel transitions from an inward-open state to the outward-open state that we observed. The notion of sequential folding supports the “BamA-elongation” model proposed by Schiffrin *et al* (2017), except that our OS-/IO-substate structures raise the possibility that β-sheet elongation and OM integration occur simultaneously. Independent of the role of BamA in OMP assembly, our finding that the essential but enigmatic BamD subunit can interact with the turns of already integrated but incompletely folded β-barrels is very striking. It is plausible that BamD supports the assembly process by sensing the extent of substrate folding or by facilitating β-strand transfer.

Finally, our work yielded significant insights into the energetics of OMP assembly. First, we obtained the first direct experimental evidence that the C-terminal side of the BamA β-barrel can modify the thickness of the membrane and therefore lower the energy requirements for the membrane integration of OMPs. Our results are in line with molecular dynamics simulations (Liu and Gumbart, 2020; Noinaj et al., 2013), *in vitro* studies that indicate that membrane thickness acts as a major barrier to OMP integration (Kleinschmidt and Tamm, 2002; Schiffrin et al., 2017), and that membrane defects accelerate β-barrel folding (Danoff and Fleming, 2015). Second, we obtained evidence that OMP assembly is not only driven by the free energy of folding but that the late stages of OMP folding are accelerated by BAM harnessing OM tension as a source of potential energy. Our experiments were inspired by an effort to explain the purpose of the unprecedented structures of the deflected EspP open β-sheets bound to BamA in an outward-open conformation. We proposed that the outward-open conformer of BamA holds the β-signal of the folding OMP at an angle at which the intrinsic structure of the open β-sheet state causes the hydrophobic transmembrane portions and aromatic girdles to deflect the OM. However, the intrinsic tension in the OM would counter this deflection and thereby forces the β-sheet to close into a β-barrel. Consistent with our model, we demonstrated that the rate of folding after the formation of a hybrid-barrel can be transiently slowed by conditions that relaxed the OM and can be accelerated when those conditions were reversed to increase the OM tension. Furthermore, because the concentration of OMPs in the OM contributes to its rigidity (Lessen et al., 2018; Rojas et al., 2018), it is plausible that the mysterious “OMP-islands” (pockets in the bacterial OM with dense OMP packing and low diffusion) generate local zones of high OM stiffness that promote the high β-barrel assembly activity attributed to them (Gunasinghe et al., 2018; Rassam et al., 2015; Ursell et al., 2012). From a different perspective, a rigid membrane might inhibit the integration of α-helical proteins which typically fold into fluid membranes. Given that transmembrane β-barrel folding occurs in an environment that is devoid of known external energy sources (e.g., ATP, GTP, or useful electrochemical gradients), the ability of BAM to catalyze transmembrane β-barrel folding by a radically different mechanism that harnesses the unusual properties of the OM might help to explain why the bacterial OM is populated almost exclusively by β-barrel proteins.

## MATERIALS AND METHODS

### Plasmids, bacterial strains, and growth media

The *E. coli* B strain BL21(DE3) (Invitrogen catalog number C600003) was used for all experiments and *E. coli* K-12 strains XL1-Blue (Agilent catalog number 200236) or NEB5α (NEB catalog number C2987H) were routinely used for cloning and mutagenesis. Strains were grown in Lysogeny Broth (LB) (Miller or Lenox formulation as indicated) supplemented with ampicillin (100 μg mL^-1^) and/or trimethoprim (50 μg mL^-1^) as necessary. Oligonucleotides and plasmids used in this study are listed in Table S1. When necessary, BAM (^His^BamABCDE) was expressed from an IPTG inducible promoter in plasmid pMTD372 and ^MBP-76^EspP was expressed from a *P*rhaB inducible promoter in plasmid pMTD607 (Doyle and Bernstein, 2019). Plasmids expressing cysteine substitution mutant derivatives of pMTD372 and pMTD607 were generated using the Q5 Site-Directed Mutagenesis Kit (NEB catalog number E0554S).

### Purification of BAM-^MBP-76^EspP native nanodiscs

*E. coli* strain BL21(DE3) transformed with plasmids expressing ^His^BamA_S425C_BCDE and ^MBP-76^EspP_S1299C_ were grown overnight in LB (Miller) at 25 °C. Overnight cultures were then washed and resuspended in fresh LB (1 culture volume) before inoculating 16 Thomson Ultra Yield flasks (each containing 1 L of LB (Miller)) at a starting OD_600_ of 0.05. Cultures were grown for 4 h (25 °C, 250 rpm), induced with 0.4 mM IPTG for 1 h, and then induced for a further 45 min with 0.2% L-rhamnose. Each culture was pelleted (5,000 x *g*, 10 min, 4 °C), resuspended in 50 mL ice-cold phosphate buffered saline (PBS; 9 g L^-1^ NaCl, 0.144 g L^-1^ KH_2_PO_4_, 0.795 g L^-1^ Na_2_HPO_4_, pH 7.4), and transferred to an Erlenmeyer flask on ice. Bacteria in each flask were treated with a final concentration of 0.4 mM 4-DPS (4,4′-dipyridyl disulfide; a thiol-specific disulfide oxidizing catalyst) for 30 min with orbital shaking at 100 rpm in packed ice, pelleted (4,500 x *g*, 10 min, 4 °C), resuspended in 25 mL ice-cold PBS containing SigmaFast EDTA free protease inhibitors (PI), and then frozen in liquid nitrogen. All 400 mL of harvested bacteria were thawed and then lysed with a Constant Systems Cell Disruptor (15,000 psi, cooled to 5 °C). Cell debris was removed (20,000 x *g*, 15 min, 4 °C) and then the lysate was ultracentrifuged (194,903 x *g*, 2 h, 4 °C) to harvest membrane pellets. Using a Dounce homogenizer, membranes were homogenized in 55 mL native-nanodisc buffer (3 % Xiran SL30010P20 (Orbiscope), 50 mM TrisHCl, 500 mM NaCl, 10 % glycerol, 1 mM EDTA, pH 8) containing freshly added PI and incubated at 4 °C for 5 h with constant inversion. The solution was ultracentrifuged (265,455 x *g*, 40 min, 4 °C) and then the supernatant was collected and diluted 1:2 with a buffer (50 mM TrisHCl, 500 mM NaCl, 10 % glycerol, pH 8 at 4 °C) containing freshly added PI. The diluted protein solution was then incubated with 25 mL StrepTactin XT superflow resin (IBA GmbH) at 4 °C overnight with constant inversion. The protein-resin solution was then transferred to a gravity column (all column steps mentioned hereafter were conducted at 4 °C) and the protein flow-through was passed over the resin a second time. The resin was washed with 10 x 50 mL TN buffer (50 mM TrisHCl, 500 mM NaCl, pH 8) at 4 °C before BAM-^MBP-76^EspP native nanodiscs were eluted with 150 mL biotin buffer (50 mM biotin, 50 mM TrisHCl, 500 mM NaCl, pH 8) at 4 °C. To concentrate and further purify the sample, imidazole (20 mM final) was added to the eluted protein which was subsequently incubated with 5 mL NiNTA resin (Qiagen) at 4 °C overnight with constant inversion. The protein solution was then transferred to a gravity column and the protein flow-through was passed over the resin twice more. The resin was washed with 3 x 10 mL of a buffer (20 mM imidazole, 50 mM TrisHCl, 500 mM NaCl pH 8) at 4 °C before BAM-^MBP-76^EspP native nanodiscs were eluted with 15 mL of elution buffer (500 mM imidazole, 50 mM TrisHCl, 150 mM NaCl pH 8) at 4 °C. The eluted protein was desalted and exchanged into TN^low^ buffer (50 mM TrisHCl, 150 mM NaCl, pH 8 at 4 °C) using Sephadex G-25 PD-10 desalting columns (Cytiva) following the manufacturers protocol before concentrating to a volume of 20-50 µL using an Amicon Ultra 0.5 mL concentrator (10 kDa cut-off). BAM-^MBP-76^EspP native nanodiscs were used immediately in grid-preparations. For each preparation, correct folding was confirmed by heat-modifiability/mobility-shift assays and activity was assessed by *in vitro* assembly-restart assays (see below).

### Cryo-EM sample preparation and imaging

BAM-^MBP-76^EspP native nanodiscs were diluted in TN^low^ buffer at a concentration of ∼2 – 8 mg L^-1^, and 3 µL of sample was applied onto glow-discharged C-flat grids (EMS CF-1.2/1.3-4Au-1. 50) for 3 sec before plunge freezing in liquid ethane using a Leica EM Grid Plunger (Leica Microsystems). Datasets were collected at the NIH Multi-Institute Cryo-EM Facility (MICEF) using a Titan Krios G3 microscope (Thermo-Fisher) operating at 300 kV. During 4 collection sessions (Figure S5, dataset 1) micrographs were collected at a magnification of 130,000x (calibrated pixel size 0.5371 Å, nominal defocus range 0.6 to 1.8 µm, 40 frames, and 60 e^-^/Å^2^ electron exposure per movie) using a Gatan K2 Summit direct electron detection camera equipped with a Gatan Quantum LS imaging energy filter with slit width set to 20 eV. After the microscope was upgraded with a Gatan K3 camera an additional collection session (Figure S5, dataset 2) was conducted at a magnification of 105,000x (calibrated pixel size 0.4281 Å, nominal defocus range 0.6 to 1.8 µm, 23 frames, and 60 e^-^/Å^2^ electron exposure per movie).

### Cryo-EM image processing

Movie frames of BAM-^MBP-76^EspP cryo-electron micrographs were motion corrected and dose-weighted with MotionCor2 in RELION 3.1 (Zheng et al., 2017; Zivanov et al., 2018). CTF estimation was determined in RELION 3.1 using Ctffind4 (Rohou and Grigorieff, 2015). Initial particle picking was done with the Laplacian-of-Gaussian-based autopicking. Picked particles were processed to generate an initial 3D reference for autopicking in RELION 3.1. A total of 25,393,510, particles, from dataset 1 collected on the K2 camera, and 9,873,900 particles, from dataset 2 on the K3 camera, were picked. Following one round of 2D classification and three rounds of 3D classification 3,996,756 particles from both datasets were merged with pixel size of 1.07 Å /pixel. Because we aimed to visualize intermediate folding states of EspP, we performed focused classification 3D classification on the 3,996,756 million particles after signal subtraction of heterogenous BamA P3, BamB, and BamD N-terminus components, which yielded a subset of 1,187,709 particles. These particles produced a 4.4 Å map using RELION 3.1. Following CTF refinement and particle polishing, the 1,187,709 particles were processed by three strategies in parallel using RELION 3.1 (Figure S5). Strategy 1 generated a 4.2 Å map after 3D refinement. Strategy 2 used 3D classification of the 1,187,709 particles in RELION 3.1 to reveal six classes of the BAM-^MBP-76^EspP complex. Strategy 3 used focused classification and refinement after signal subtraction of BamA P3, BamB, and BamD N-terminus revealed three folding states of EspP. Particles from strategy 1, the six classes in strategy 2, and the 3 states in strategy 3 were moved from the RELION 3.1 pipeline to cryoSPARC for further cryo-EM image processing (Punjani et al., 2017; Punjani et al., 2020). Following pruning of the particle sets by rounds of heterogenous refinement and final refinements, the following cryo-EM maps were obtained: (1) a 3.6 Å map of BAM-^MBP-76^EspP from strategy 1; (2) six cryo-EM maps capturing the motion in the soluble subunits of BAM-^MBP-76^EspP from strategy 2 [class 1 (4.5 Å), class 2 (4.3 Å), class 3 (4.2 Å), class 4 (4.3 Å), class 5 (4.3 Å), and class 6 (4.2 Å)]; (3) three cryo-EM maps following focused classification/refinement of the substrate region produced the OS-state (4.3 Å), IO-state (4.3 Å), and the B/CO-state (4.8 Å) stemming from strategy 3. Local resolution filtered maps were produced in cryoSPARC.

### Model building and refinement

Initial fitting of BAM-^MBP-76^EspP subunits into cryo-EM maps was done manually in UCSF Chimera (Pettersen et al., 2004) using Bam complex subunits from PDB 5D0O (Gu et al., 2016), EspP from PDB 2QOM and 3SLJ (Barnard et al., 2007; Barnard et al., 2012), and lipopolysaccharide from 5W7B (Gorelik et al., 2018). For the high-resolution 3.6 Å map, manual building/corrections of BamA subunits, EspP and LPS was done in Coot 0.9 and Isolde followed by model refinement using Rosetta and real-space refinement in Phenix (Adams et al., 2010; Croll, 2018; Emsley et al., 2010; Wang et al., 2016). The high-resolution atomic model derived from the 3.6 Å map was used as a starting model for building models of the six classes (from strategy 2) and the three focused states (from strategy 3). Because the substrate region in the focused maps is observed at low resolution, some of the EspP β-barrel N-terminus was docked into the map. The membrane interacting regions of EspP were better defined, could be identified by the orientations of proteins in OPM server (Lomize et al., 2012), and modeled into the cryo-EM map using Rosetta, Coot and Isolde (Croll, 2018; Emsley et al., 2010; Wang et al., 2016). The cryo-EM data collection, final refinement, and validation statistics for the 10 atomic models are presented in Table S2. Structural analysis, measurements and figures were prepared in Chimera and ChimeraX (Pettersen et al., 2021).

### Data and code availability

Structural data supporting findings in this study have been deposited in the Protein Data Bank (PDB) and the Electron Microscopy Data Bank (EMDB). The accession codes of the cryo-EM maps and accompanying atomic models have been provided for: (1) BAM-^MBP-76^EspP *high-resolution* (EMDB-xxxxx, PDB:xxx): (2) BAM-^MBP-76^EspP *class 1* (EMDB-xxxxx, PDB:xxx): (3) BAM-^MBP-76^EspP *class 2* (EMDB-xxxxx, PDB:xxx): (4) BAM-^MBP-76^EspP *class 3* (EMDB-xxxxx, PDB:xxx): (5) BAM-^MBP-76^EspP *class 4* (EMDB-xxxxx, PDB:xxx): (6) BAM-^MBP-76^EspP *class 5* (EMDB-xxxxx, PDB:xxx): (7) BAM-^MBP-76^EspP *class 6* (EMDB-xxxxx, PDB:xxx): (8) BAM-^MBP-76^EspP *open-sheet EspP state* (EMDB-xxxxx, PDB:xxx): (9) BAM-^MBP-76^EspP *intermediate-open EspP state* (EMDB-xxxxx, PDB:xxx): (10) BAM-^MBP-76^EspP *barrelized EspP/ continuous open BamA state* (EMDB-xxxxx, PDB:xxx).

### In vivo disulfide-bond formation assay

To observe site-specific interactions between BamD and the EspP β-barrel *in vivo*, disulfide-bond formation assays were conducted essentially as described (Doyle and Bernstein, 2019, 2021). Briefly, strains containing appropriate plasmids were grown overnight from a single colony in 10 mL LB (Miller) at 25 °C with orbital shaking (250 rpm). Cultures were pelleted (3000 x g, 5 min, 4 °C), washed with 10 mL LB (Miller), and resuspended in 10 mL LB (Miller) before inoculating 10 mL LB (Miller) subcultures at OD_600_ = 0.05. After cultures were grown for 4 h (25 °C, 250 rpm) to OD_600_ ∼0.4 - 0.6, a final concentration of 0.4 mM IPTG was added to induce expression of BAM for 1 h. Subsequently, a final concentration of 0.2% L-rhamnose was added to induce expression of ^MBP-76^EspP for 45 min. 1 mL samples were aliquoted into tubes on ice, pelleted (10,000 x g, 2 min, 4 °C), resuspended in 1 mL of ice-cold PBS, and incubated on ice with 4-DPS at a concentration of 0.2 mM (or an equivalent volume of ethanol for mock treatment controls). After 30 min, samples were pelleted (10,000 x g, 2 min, 4 °C) and resuspended in 0.5 mL ice-cold PBS. Bacteria were then lysed, and proteins were precipitated by adding a final concentration of 10% (v/v) trichloroacetic acid (TCA) and 4 mM phenylmethanesulfonyl fluoride (PMSF) and incubating for 10 min on ice. The precipitated proteins were pelleted (20,817 x g, 10 min, 4 °C), washed with 0.6 mL ice-cold acetone, re-pelleted, and air-dried at 37 °C for 20 min. Proteins were resuspended in 2x SDS protein gel loading solution (Quality Biological) in a volume normalized to an OD_600_ measurement recorded immediately as subculture samples were taken (volume in μL = 200 x OD_600_). Samples were heated to 99 °C for 15 min and aliquots (5 µL) resolved by SDS-PAGE on 8 % – 16 % Tris-glycine minigels (Invitrogen) (150 V, 1 h 47 min, room temperature) before being transferred to nitrocellulose for immunoblot analysis.

### Heat-modifiability/gel mobility-shift assay

To observe the folded states of the BamA-EspP hybrid-barrel, purified BAM-^MBP-76^EspP native nanodiscs were diluted 1:9 in ice-cold TN buffer before aliquots were further diluted 1:9 in modified loading buffer (2x SDS protein gel loading solution serially diluted 1:1 with 20 % glycerol twice and then 1:1 again with TN buffer for a final SDS concentration of 0.5%) on ice. Aliquots were either heated to 99 °C for 10 min or retained on ice and proteins were immediately resolved by cold-SDS-PAGE (gel tank in packed ice, running at 150 V, 4 °C cold room). Gels were subsequently Coomassie Brilliant Blue (R-250) stained to detect proteins.

### In vivo ^MBP-76^EspP assembly-restart assays

To monitor the final stages of assembly of EspP after the formation of a hybrid-barrel intermediate with BamA, bacteria containing plasmids that express BAM and ^MBP-76^EspP were cultured overnight from a single colony in 10 mL LB (Lenox) at 25 °C with orbital shaking (250 rpm). Cultures were pelleted (4,000 x g, 3 min, 4 °C), washed with LB (Lenox), and resuspended in 10 mL LB (Lenox) before inoculating 10 mL LB (Lenox) subcultures at OD_600_ = 0.05. To create a pool of molecules at a hybrid-barrel intermediate stage of assembly in bacteria, subcultures were grown and the expression of BAM and ^MBP-76^EspP was induced as in the disulfide-bond formation assays described above. Aliquots (1 mL samples) were then pelleted (10,000 x g, 2 min, 20 °C), resuspended in equivalent volumes of either LB (Lenox) or LB (Lenox) containing 0.8 M sorbitol (LB-Sorbitol), and pre-incubated in a Thermomixer (Eppendorf) (20 °C, 350 rpm). After 5 min pre-incubation, 200 μg mL^−1^ PK (or an equivalent volume of 50 mM TrisHCl pH 8 for mock-treated controls) was added and bacteria were incubated (20 °C, 350 rpm) for 0, 2, 5, 10, and 20 min. For experiments requiring media exchange from LB-Sorbitol, samples incubated with PK for 5 min were pelleted (15,000 x g, 20 s, 20 °C), resuspended in 1 mL of either LB (Lenox) or LB-Sorbitol (media pre-equilibrated to 20 °C), and further incubated (20 °C, 350 rpm) until 10 or 20 min after PK addition. For experiments requiring disulfide-crosslinking, bacteria incubated with PK for 5 min were pelleted (15,000 x g, 20 s, 4 °C), resuspended on ice in 1 mL ice-cold LB (Lenox) or LB-Sorbitol (matching the previous incubation medium for each sample), and incubated on ice for 2 min in the presence of 4-DPS (0.2 mM final concentration). To stop reactions at required time-points, bacteria were pelleted (15,000 x g, 20 s, 4 °C), resuspended in 0.5 mL ice-cold LB (Lenox) or LB-Sorbitol (matching the previous incubation medium for each sample), and TCA precipitated as described above. Precipitated proteins were solubilized and resolved by SDS-PAGE as described above.

### Immunoblotting and image quantitation

The iBlotII transfer device (Life Technologies) was routinely used to transfer protein gels to nitrocellulose membranes. Immunoblotting buffer [Odyssey Blocking Buffer (Li-Cor) and PBS (mixed together at a 1:1 ratio)] supplemented with 0.01 % Tween-20 was used for blocking steps and as a diluent for primary and secondary antibodies. Monoclonal mouse anti-StrepII and anti-His antibodies were obtained from QIAGEN (catalog number 34850) and Genscript (catalog number A00186), respectively. Polyclonal rabbit anti-BamD and anti-EspP_βC_ have been described previously (Pavlova et al., 2013). Infra-red Goat anti-mouse Ig secondary antibodies (anti-mouse 800CW IRDye, catalog number 926-32210) or anti-rabbit Ig (anti-rabbit 680LT IRDye, catalog number 926-680210) were obtained from Li-Cor. Membranes were blocked overnight, incubated with primary antibodies for 18 h, washed twice with PBS-T (PBS supplemented with 0.01 % Tween-20), incubated for 2 h with secondary antibodies, and washed twice with PBS-T and three times with PBS before air drying (37 °C, 20 min). Dried membranes were scanned using maximum quality and resolution settings with an Amersham Typhoon 5 imager (GE Healthcare) outfitted with 785 nm and 685 nm lasers and IRlong 825BP30 and IRShort 720BP20 filters. Within-lane pixel intensities were measured using Fiji software (v2.0.0-rc-68/1.52 g) and used to calculate the fraction of the band of interest relative to other bands of interest [e.g. for assembly restart assays, the fraction of folded EspP β-barrel was determined by using the formula [folded EspP β-barrel/sum of EspP signals)].

## ACKNOWLEDGEMENTS

This work was supported by the National Institute of Diabetes and Digestive and Kidney Diseases Intramural Research Program. JRJ is a recipient of NIGMS MOSAIC K99/R00 and NIDDK Nancy Nossal awards. The structural studies were performed at the NIH Multi-Institute Cryo-EM Facility (MICEF) and the NIDDK Cryo-EM core facility and utilized computational resources from the NIH HPC Biowulf cluster (http://hpc.nih.gov). We thank Huaibin Wang, Haifeng He, and Bertram Canagarajah for technical support with electron microscopy and computing.

## AUTHOR CONTRIBUTIONS

The study was originally conceived by MTD and HDB, but all authors contributed to experimental design. The experiments and data processing were conducted by MTD and JRJ. The paper was written and edited by all authors. The project was supervised by JEH and HDB.

