## Supplemental Data for "Cryo-EM structures reveal multiple stages of bacterial outer membrane protein folding"

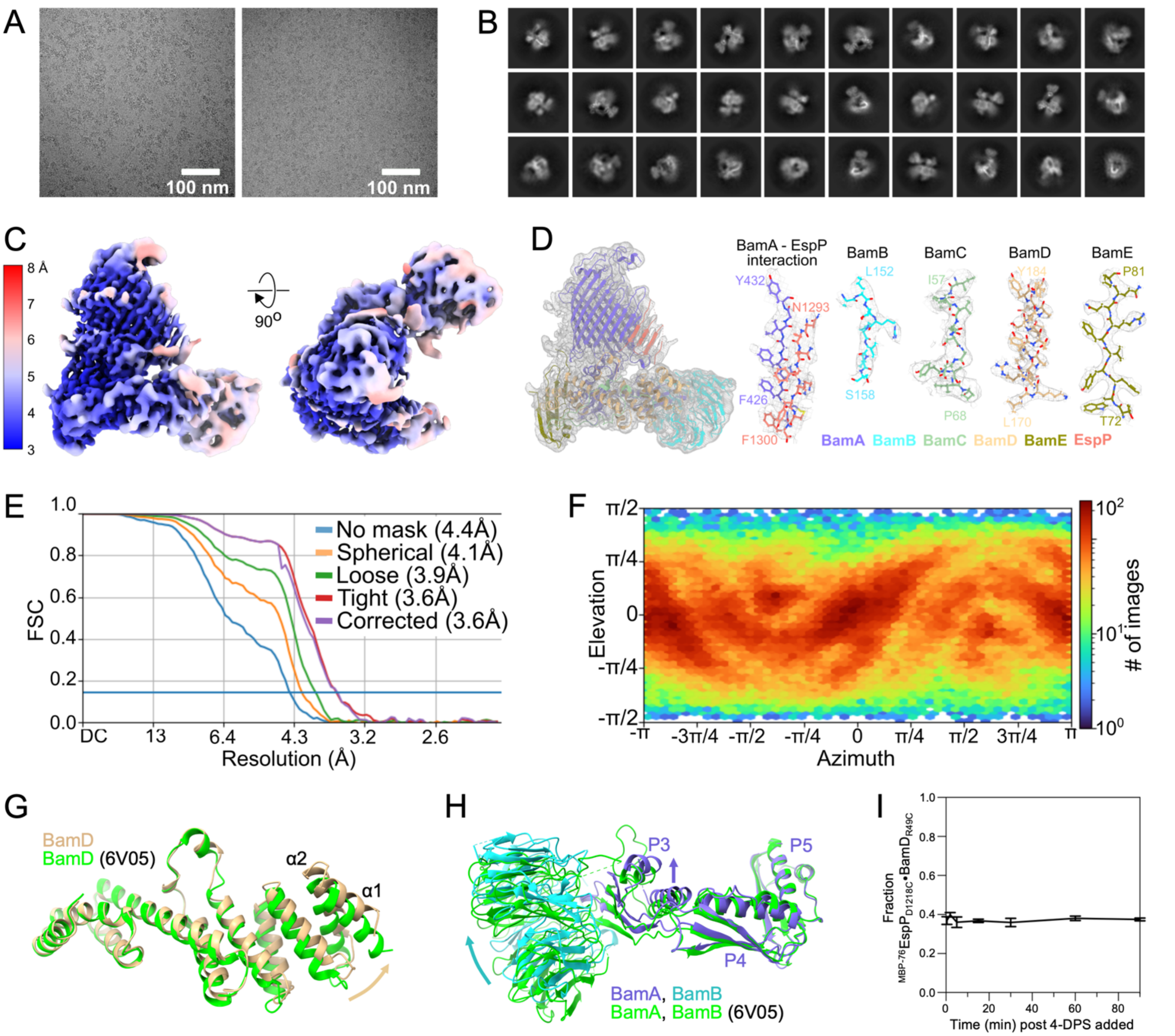


**Supplementary Figure 1: Generation and validation of BAM-$^{MBP-76}$EspP high-resolution map and atomic model, in relation to Figure 1.**

**(A)** Representative cryo-EM micrographs of BAM-$^{MBP-76}$EspP OM-nanodiscs. **(B)** Representative 2D classes. **(C)** Map colored by local resolution. **(D)** Left, BAM-$^{MBP-76}$EspP atomic model in the map. Right, representative fits of BamA-EspP β-barrel hybridization interface and BamB-E models to the map. **(E)** Gold-standard Fourier shell correlation (FSC). **(F)** Orientation distribution of particles used in reconstruction. **(G)** BAM-BamA$_{\Delta L1}$ hybrid-barrel (6V05) (green) aligned with the BAM-$^{MBP-76}$EspP model (on BamA POTRA5 residues Y348 – R421). Only BamD is shown (colors as in Figure 1). Compared to BAM-BamA$_{\Delta L1}$, the N-terminal α-helices of the BamD subunit in the BAM-$^{MBP-76}$EspP structure expands towards the membrane. **(H)** As in **G** except showing only BamA POTRA domains P3-5 and BamB. For BAM-$^{MBP-76}$EspP, P3 and BamB are shifted towards the membrane. **(I)** *E. coli* BL21(DE3) expressed $^{His}$BamABCD$_{R49C}$E and $^{MBP-76}$EspP$_{D1218C}$ as in Figure 1E, except that samples were taken at multiple time points after the addition of the 4-DPS oxidant (±SEM, n = 3).

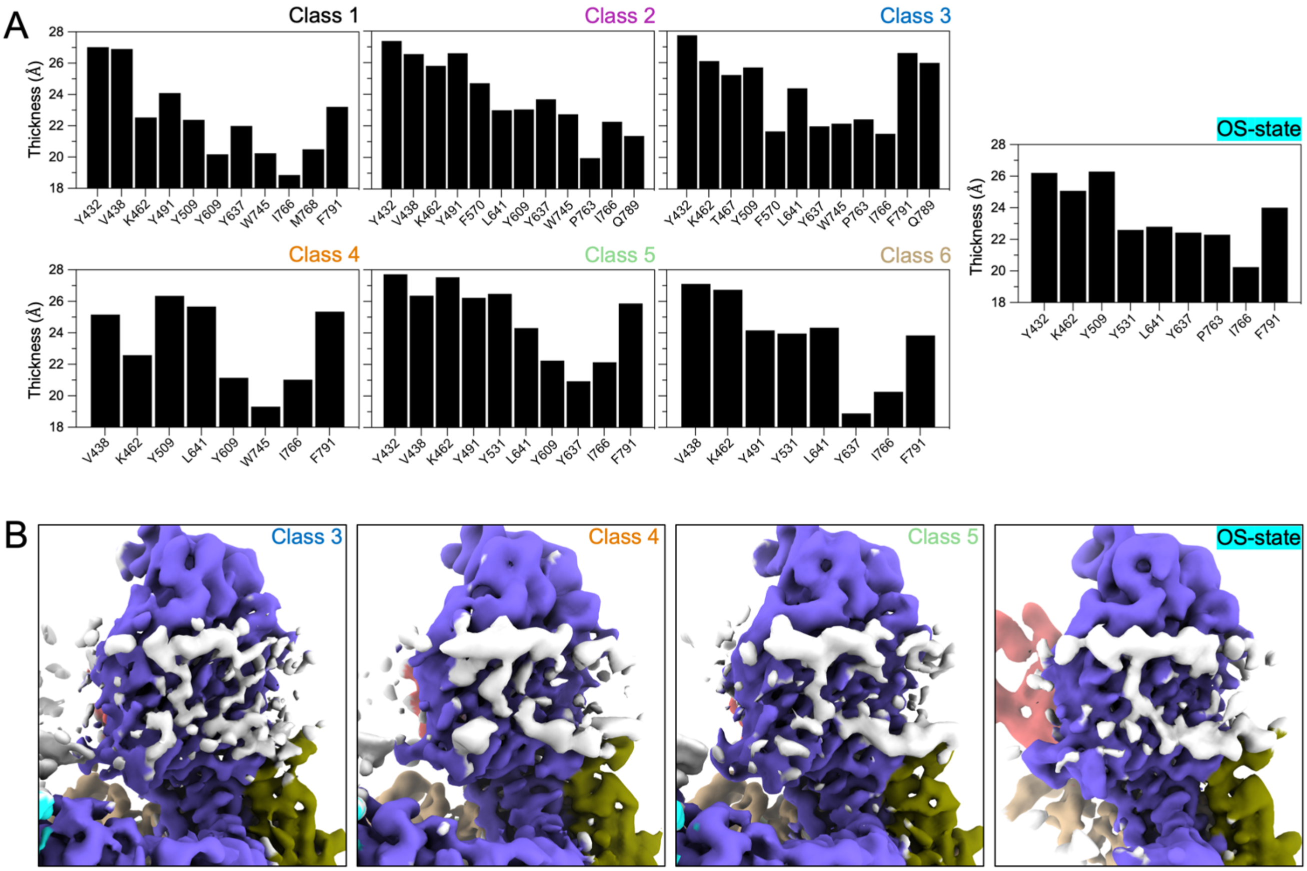


**Supplementary Figure 2: OM thinning and lipid/LPS stabilization during OMP assembly is apparent in multiple different maps, in relation to Figure 2.**

**(A)** The thicknesses of membrane density was measured surrounding the BamA β-barrel at positions close to indicated residues for the other generated maps that are not shown in Figure 2A (see Figures 3, 4 & S3). **(B)** Views of selected maps with proteins colored as in Figure 1 and high-density stabilized membrane components colored in white. Density consistent with lipid A head groups and a stabilized acyl chain in the high-resolution map in Figure 2 is also reproduced in independently derived maps.

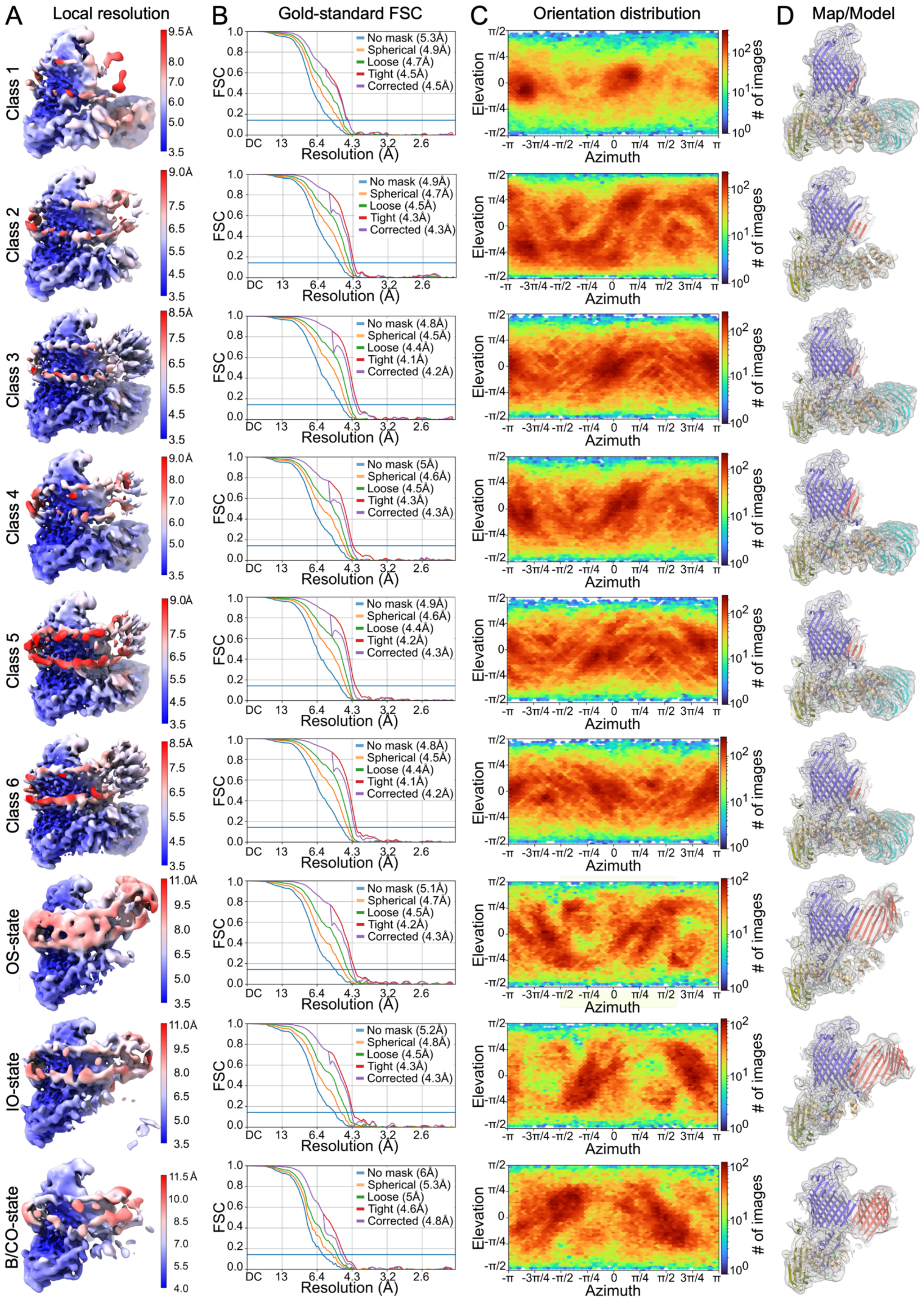


**Supplementary Figure 3 (legend next page)**

**Supplementary Figure 3: Validation of BAM-$^{MBP-76}$EspP whole-particle maps (classes 1-6) and focused maps, relating to Figures 3 and 4.**

**(A)** Column shows each reconstructed cryo-EM map colored by local resolution. **(B)** Column shows gold-standard Fourier shell correlation (FSC) curves for each map. **(C)** Column shows orientation distributions of particles used in each reconstruction. **(D)** Column shows refined BAM-$^{MBP-76}$EspP atomic models within each map.

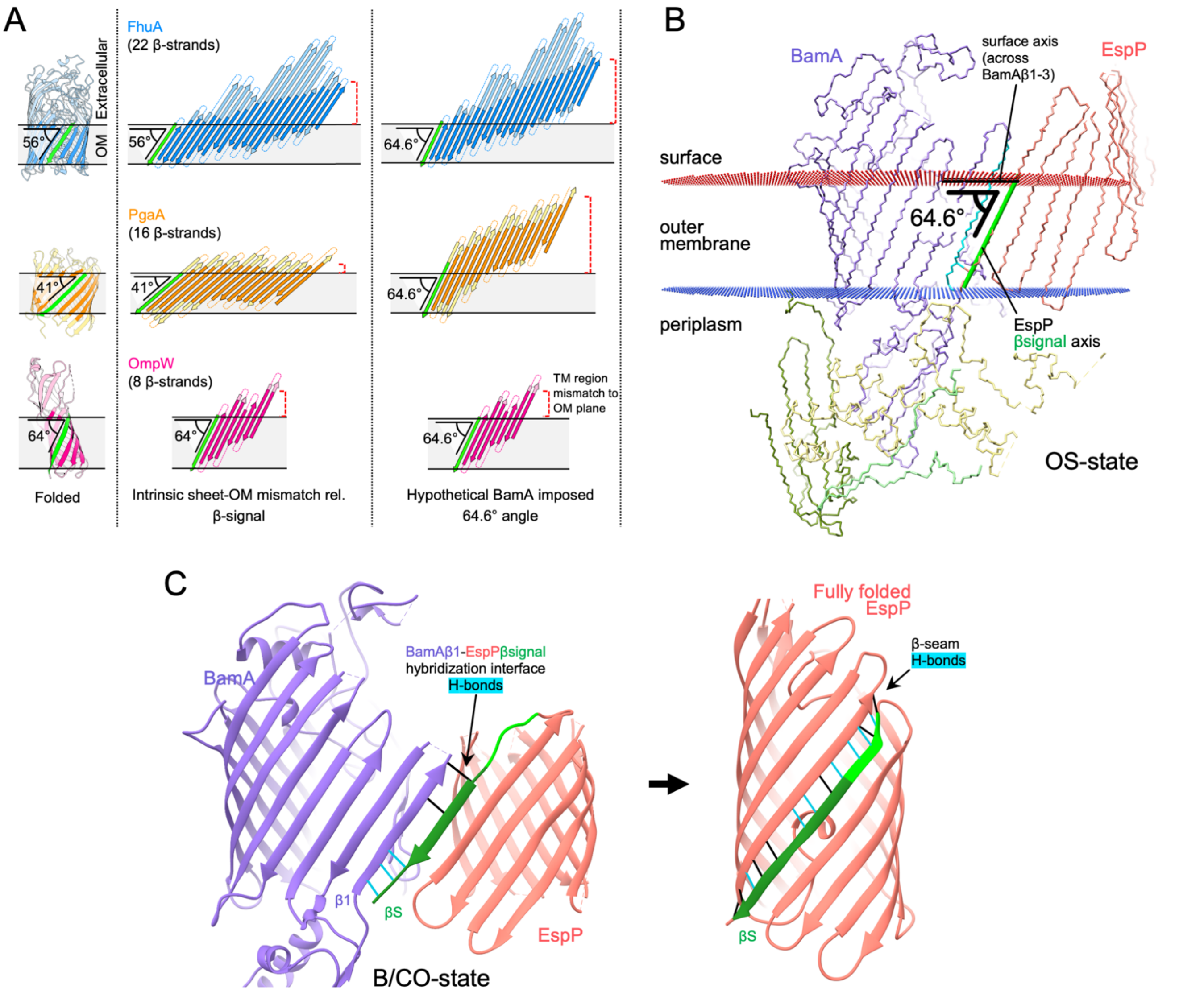


**Supplementary Figure 4: BamA accentuated angular antagonism of incompletely folded β-barrels and relative instability of the hybrid-barrel state, relating to Figure 5.**
**(A)** *E. coli* β-barrels of varied sizes (FhuA; PDB 1FCP, PgaA; PDB 4Y25, OmpW; PDB 2F1V) (left column) were hypothetically unraveled (middle column) with the position of the conserved β-signal (green) held constant to its orientation when fully folded showing intrinsic mismatch between the membrane plane and the transmembrane portion of β-strands (dark shade of each strand). Right column shows expanded β-sheets at a hypothetical BamA imposed angle (extrapolated from measured angle of EspP β-signal when bound to BamA). We speculate that BamA holds OMPs at a high angle to harness OM tensile forces to accelerate β-barrel folding. **(B)** The BAM-$^{\text{MBP-76}}$EspP OS-state model shown with membrane plane based on the BAM subunits [OPM server (Lomize et al., 2012)]. The β-signal axis (green) was calculated with the membrane spanning backbone. The surface axis (black) was calculated across BamA β-strands 1-3. The angle between the surface and β-signal axes is 64.6°. **(C)** The hybridization interface of the BAM-$^{\text{MBP-76}}$EspP B/CO-state has fewer hydrogen (H)-bonds than when the β-seam when EspP (2QOM) is fully folded. H-bonds were identified using Chimera FindHBond that fit the precise criteria (cyan) or with relaxed tolerances (0.4 Å / 20.0°) (black). Dark green = 1292-1300.

### Supplementary Figure 5 (legend next page)

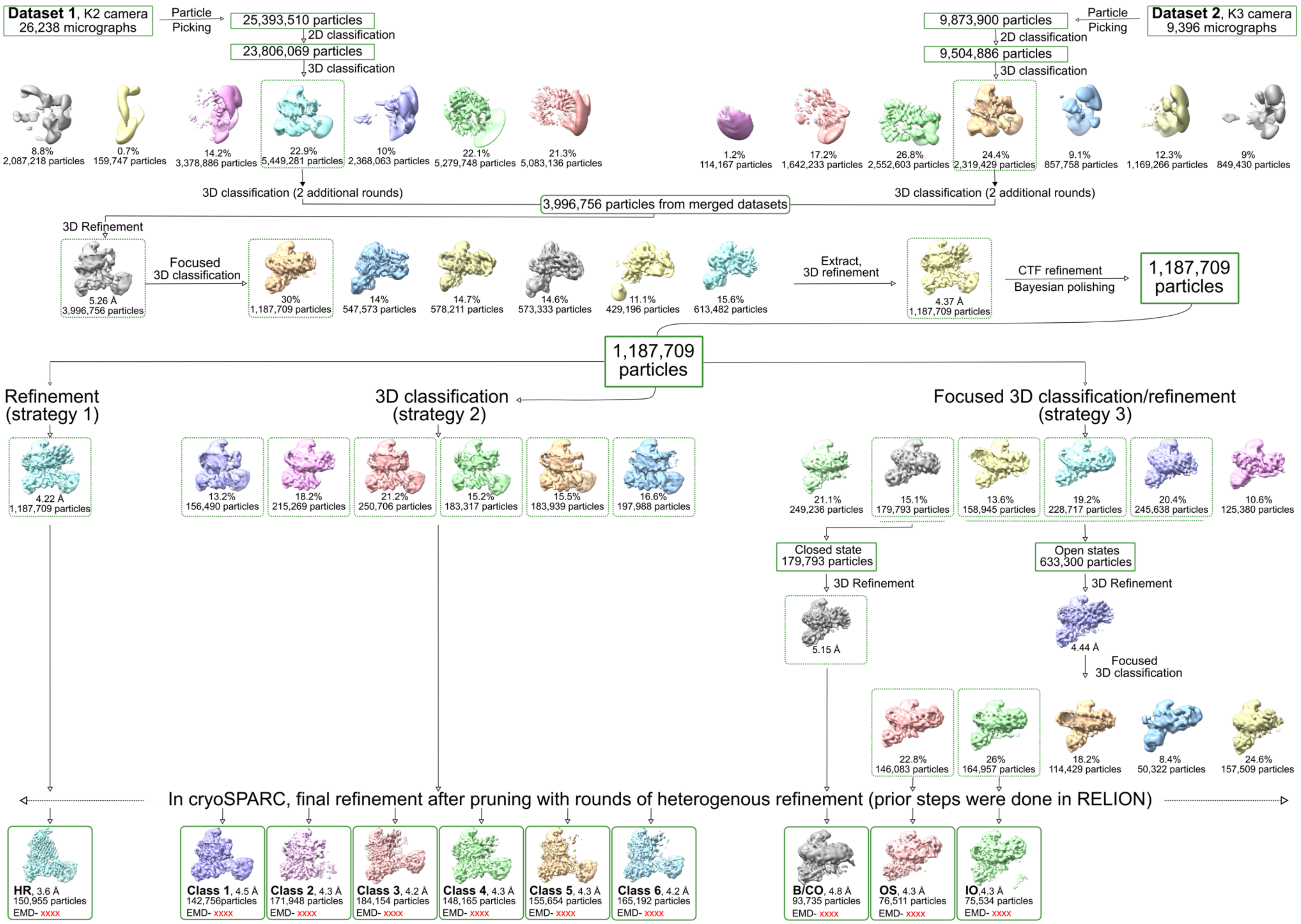

**Supplementary Figure 5: Flowchart of BAM-$^{MBP-76}$EspP Cryo-EM reconstructions.**
The flowchart outlines the processing approaches that led to the high-resolution (**HR**) map, the six whole particle classes (**Classes 1- 6**) and the open-sheet (**OS**), intermediate-open (**IO**) and barrelized/continuous open (**B/CO**) focused reconstructions.

| ssDNA oligo | Sequence | Notes |
|---|---|---|
| mtd388 | `TGCTGGAAAGTGACAGCCCGTG` | F, primer for *espP D1218C* substitution |
| mtd389 | `CTTACCAGAGAAGGATTTACCCACATC` | R, primer for *espP D1218C* substitution |
| mtd390 | `TGCCAGGCAATAACGCAACTGGAAG` | F, primer for BamD *R49C* substitution |
| mtd391 | `CCAGTTACCGTCCTGCAGC` | R, primer for BamD *R49C* substitution |
| mtd392 | `TGCCAAGGGTTCTTTGGCGTCGATC` | F, primer for BamD *L124C* substitution |
| mtd393 | `CGCACTGTCATCCAGCGC` | R, primer for BamD *L124C* substitution |

| Plasmid | Notes | Source |
|---|---|---|
| pMTD372 | pTrc99a::$^{His8}$*bamABCDE*, $P_{Trc}$, Amp$^{R}$ | 1 |
| pMTD607 | 'pRha$^{MBP-76}$EspP' (pSCrhaB2::*espP*$_{SS}$*-TS-malE-espP*$_{948-984}$*-tev-espP*$_{985-1300}$), $P_{rhaB}$, Tmp$^{R}$ | 1 |
| pMTD710 | pTrc99a::$^{His8}$*bamA*$_{S425C}$*-BCDE* | 1 |
| pMTD712 | pRha$^{MBP-76}$EspP$_{S1299C}$ | 1 |
| pMTD1582 | pRha$^{MBP-76}$EspP$_{D1218C}$ (*espP*$_{D1218C}$ substitution on pMTD607 with mtd388/389 primers) | This work |
| pMTD1583 | pTrc99a::$^{His8}$*bamABC bamD*$_{R49C}$ *bamE* (*bamD*$_{R49C}$ substitution on pMTD372 with mtd390/391 primers) | This work |
| pMTD1593 | pTrc99a::$^{His8}$*bamABC bamD*$_{L124C}$ *bamE* (*bamD*$_{L124C}$ substitution on pMTD372 with mtd392/393 primers) | This work |

**Supplementary Table 1: Oligonucleotides and plasmids used in this study.**
Abbreviations: SS = signal sequence, *malE* = *malE* codons *26-392*, MBP = maltose binding protein (26-392), Amp$^{R}$ = Ampicillin resistance, Tmp$^{R}$ = Trimethoprim resistance, TS = TwinStrepII tag, His8 = His x 8 tag.

(1) Doyle, M.T., and Bernstein, H.D. (2019) Bacterial outer membrane proteins assemble via asymmetric interactions with the BamA β-barrel. Nat Commun 10, 3358.

| | Bam-EspP HR state EMDB-xxxx PDB xxxx | Bam-EspP class 1 EMDB-xxxx PDB xxxx | Bam-EspP class 2 EMDB-xxxx PDB xxxx | Bam-EspP class 3 EMDB-xxxx PDB xxxx | Bam-EspP class 4 EMDB-xxxx PDB xxxx | Bam-EspP class 5 EMDB-xxxx PDB xxxx | Bam-EspP class 6 EMDB-xxxx PDB xxxx | Bam-EspP OS state EMDB-xxxx PDB xxxx | Bam-EspP IO state EMDB-xxxx PDB xxxx | Bam-EspP B/CO state EMDB-xxxx PDB xxxx |
|---|---|---|---|---|---|---|---|---|---|---|
| **Data collection & processing** | | | | | | | | | | |
| Magnification, K2 dataset | 130,000 | 130,000 | 130,000 | 130,000 | 130,000 | 130,000 | 130,000 | 130,000 | 130,000 | 130,000 |
| Magnification, K3 dataset | 105,000 | 105,000 | 105,000 | 105,000 | 105,000 | 105,000 | 105,000 | 105,000 | 105,000 | 105,000 |
| Voltage (kV) | 300 | 300 | 300 | 300 | 300 | 300 | 300 | 300 | 300 | 300 |
| Electron exposure ($e^-$/Å2 ) | 60 | 60 | 60 | 60 | 60 | 60 | 60 | 60 | 60 | 60 |
| Defocus range (μm) | 0.6 - 1.8 | 0.6 - 1.8 | 0.6 - 1.8 | 0.6 - 1.8 | 0.6 - 1.8 | 0.6 - 1.8 | 0.6 - 1.8 | 0.6 - 1.8 | 0.6 - 1.8 | 0.6 - 1.8 |
| Pixel size K2 dataset (Å) | 0.5371 | 0.5371 | 0.5371 | 0.5371 | 0.5371 | 0.5371 | 0.5371 | 0.5371 | 0.5371 | 0.5371 |
| Pixel size K3 dataset (Å) | 0.4281 | 0.4281 | 0.4281 | 0.4281 | 0.4281 | 0.4281 | 0.4281 | 0.4281 | 0.4281 | 0.4281 |
| Symmetry imposed | C1 | C1 | C1 | C1 | C1 | C1 | C1 | C1 | C1 | C1 |
| Initial particle images (no.) | 25,393,510 | 25,393,510 | 25,393,510 | 25,393,510 | 25,393,510 | 25,393,510 | 25,393,510 | 25,393,510 | 25,393,510 | 25,393,510 |
| Final particle images (no.) | 150,955 | 142,756 | 171,948 | 184,154 | 148,165 | 155,654 | 165,192 | 76,511 | 75,534 | 93,735 |
| Map resolution (Å) | 3.6 | 4.5 | 4.3 | 4.2 | 4.3 | 4.3 | 4.2 | 4.3 | 4.3 | 4.8 |
| FSC threshold | 0.143 | 0.143 | 0.143 | 0.143 | 0.143 | 0.143 | 0.143 | 0.143 | 0.143 | 0.143 |
| **Refinement** | | | | | | | | | | |
| Initial model used (PDB code) | 5D0O, 3SLJ, 5W7B | | | | | | | | | |
| Model resolution (Å) | 3.5 | 4.7 | 4.3 | 4.3 | 4.4 | 4.3 | 4.3 | 4.4 | 4.4 | 7.2 |
| FSC threshold | 0.5 | 0.5 | 0.5 | 0.5 | 0.5 | 0.5 | 0.5 | 0.5 | 0.5 | 0.5 |
| Map-model CC | 0.78 | 0.7 | 0.75 | 0.74 | 0.72 | 0.74 | 0.76 | 0.71 | 0.7 | 0.64 |
| Model composition | | | | | | | | | | |
| Non-hydrogen atoms | 10,285 | 10,061 | 7,634 | 10,385 | 10,402 | 10,473 | 10,535 | 7,318 | 7,210 | 6,842 |
| Protein residues | 1,311 | 1,287 | 962 | 1,328 | 1,330 | 1,339 | 1,347 | 928 | 915 | 869 |
| Ligands | 1 | - | - | - | - | - | - | - | - | - |
| R.M.S. deviations | | | | | | | | | | |
| Bond length (Å) | 0.005 | 0.005 | 0.004 | 0.004 | 0.004 | 0.003 | 0.003 | 0.007 | 0.005 | 0.004 |
| Bond length (°) | 0.741 | 0.896 | 0.831 | 0.909 | 0.815 | 0.891 | 0.727 | 0.855 | 0.905 | 0.86 |
| Validation | | | | | | | | | | |
| MolProbity score | 1.01 | 1.40 | 1.10 | 1.04 | 1.18 | 1.12 | 0.96 | 1.19 | 1.10 | 1.31 |
| Clashscore | 2.29 | 7.20 | 3.09 | 2.55 | 3.92 | 3.31 | 1.94 | 4.07 | 3.06 | 5.78 |
| Poor rotamers (%) | 0.00 | 0.00 | 0.00 | 0.00 | 0.00 | 0.00 | 0.00 | 0.00 | 0.00 | 0.00 |
| Ramachandran plot | | | | | | | | | | |
| Favored (%) | 98.98 | 98.72 | 98.83 | 98.62 | 98.70 | 98.93 | 98.79 | 99.10 | 98.63 | 98.54 |
| Allowed (%) | 1.02 | 1.28 | 1.17 | 1.38 | 1.30 | 1.07 | 1.21 | 0.90 | 1.37 | 1.46 |
| Disallowed (%) | 0.00 | 0.00 | 0.00 | 0.00 | 0.00 | 0.00 | 0.00 | 0.00 | 0.00 | 0.00 |

**Supplementary Table 2: Cryo-EM data collection, refinement, and validation statistics.**

**Video S1: Conformational changes of BamA POTRA domains and BAM lipoproteins during OMP folding, in relation to Figure 3.**

This video shows a transition between the BAM-$^{MBP-76}$EspP structural Classes 1 and 3-5 (except Class 2 in which BamB was not modelled). Video plays through twice in the following order: Class 5→3→4→6→1→5. Made with Chimera Morph Conformations tool after models were aligned on POTRA5 (residues Y348 – R421).

**Video S2: Conformational changes during EspP β-barrel folding observed by focused classification, in relation to Figure 4.**

This video shows a transition between the BAM-$^{MBP-76}$EspP OS, IO, and B/CO states. Video shows a side view and then a surface view of the complex and plays through twice in the following order: OS→IO→B/CO. Made with Chimera Morph Conformations tool after models were aligned on POTRA5 (residues Y348 – R421).
